# Condensate-driven interfacial forces reposition DNA loci and measure chromatin viscoelasticity

**DOI:** 10.1101/2023.02.27.530281

**Authors:** Amy R. Strom, Yoonji Kim, Hongbo Zhao, Natalia Orlovsky, Yi-Che Chang, Andrej Košmrlj, Cornelis Storm, Clifford P. Brangwynne

## Abstract

Biomolecular condensates assemble in living cells through phase separation and related phase transitions. An underappreciated feature of these dynamic molecular assemblies is that they form interfaces with cellular structures, including membranes, cytoskeleton, DNA and RNA, and other membraneless compartments. These interfaces are expected to give rise to capillary forces, but there are few ways of quantifying and harnessing these forces in living cells. Here, we introduce VECTOR (<u>V</u>isco<u>E</u>lastic <u>C</u>hromatin <u>T</u>ethering and <u>OR</u>ganization), which uses light-inducible biomolecular condensates to generate capillary forces at targeted DNA loci. VECTOR can be utilized to programmably reposition genomic loci on a timescale of seconds to minutes, quantitatively revealing local heterogeneity in the viscoelastic material properties of chromatin. These synthetic condensates are built from components that naturally form liquid-like structures in living cells, highlighting the potential role for native condensates to generate forces and do work to reorganize the genome and impact chromatin architecture.

## Introduction

Biomolecular condensates are membraneless assemblies that form within living cells through liquid-liquid phase separation (LLPS) and related phase transitions^1-2^. These include P granules^3^, T cell receptor clusters^4^, stress granules^5^, the pyrenoid^6^, and nucleoli^7^, many of which are known to physically interact with other condensates and cellular structures. Within the nucleus, some condensates are directly associated with chromatin, the nucleic acid- and protein-based polymer that stores genetic information. Chromatin-associated condensates are thought to be involved in multiple nuclear functions including coordinating transcriptional activation^8-9^ or repression^10^ of specific euchromatic sequences, and formation and silencing of heterochromatin domains^11–13^. Aberrant spatiotemporal regulation or locus targeting of chromatin-interacting condensates may be associated with diseased states^14-15^.

Foundational to the study of phase separation within the complex polymeric environment of the nucleus, decades of work in soft (non-living) matter show how the processes of nucleation, growth, and coarsening associated with phase separation can be impacted by structured environments^16-17^ also shown theoretically^18-19^. Consistent with a role for related effects in living cells, previous studies have reported on the ways in which the viscoelastic intracellular milieu impacts phase separation, for example through interactions between condensates and the cytoskeleton^20-21^, or in the nucleus through the impact of chromatin on phase separation^2,19,22^ and the nuclear F-actin network in *Xenopus laevis* oocytes^23^. In the context of the cell nucleus, condensates have been shown to favor forming in mechanically softer regions^24^, and chromatin can also impact condensate coarsening dynamics^25-26^. Thus, the material state of the cellular interior impacts the formation, positioning and organization of condensates.

One of the challenges to elucidating the interplay between condensates and intracellular structure is our incomplete current understanding of the mechanical environment inside of living cells, particularly inside the nucleus. Indeed, while the material state of chromatin influences functions including replication, transcription, and protection from DNA damage^27^, there is no consensus model describing this material state. Several studies suggest that chromatin behaves as a solid on the mesoscale^28-29^, while others have suggested a liquid-like state^30-31^. Some of these differences likely reflect the “bulk” measurements that average over multiple structures; for example, methods utilizing active mechanical response measurements apply forces to the nuclear exterior, and therefore include contributions from the nuclear lamina and envelope. By contrast, there are extremely few methods of direct force-response measurements utilizing local force application on singular chromatin loci^31^.

In considering potential means of generating and harnessing forces within living cells, it is noteworthy that interfacial (surface) tension between immiscible phases can give rise to capillary forces, which likely also manifest with condensates in living cells^7,24,32,33,34^. These mechanical forces arise from the fact that non-spherical condensates are energetically unfavorable, resulting in a thermodynamic driving force that can round up the condensate into a spherical shape to decrease free energy by minimizing surface area. Indeed, recent studies have shown that model condensate systems can exhibit interfacial forces that drive intracellular restructuring^20,21,35,36^. These studies suggest that condensates could function in part like molecular motors, applying picoNewton(pN)-level forces to cellular objects; instead of utilizing ATP hydrolysis, these forces would result from free energy stored in their interfaces. Exploiting condensate-driven interfacial interactions in the context of the nucleus, could provide a method for precisely probing local chromatin material properties, and help uncover general principles underlying genome organization and function. However, there are currently extremely few approaches for measuring and controlling interfacial forces in living cells.

Here, we introduce VECTOR (<u>V</u>isco<u>E</u>lastic <u>C</u>hromatin <u>T</u>ethering and <u>OR</u>ganization), a system that creates interfacial interactions between an inducible synthetic condensate and target chromatin loci, to apply a pulling force on the attached chromatin during condensate dissolution, resulting in their repositioning. Locus repositioning is rapid (∼2 min), efficient (66% of attached loci are repositioned), specific, and precise over micron distances. We combine analytical modeling and simulations to understand the work done through interfacial interactions and to make quantitative predictions of differential viscoelastic properties across chromatin regions. With these tools, we characterized the material state of chromatin as a viscoelastic fluid consistent with the standard linear fluid Jeffreys model^37-38^ with local heterogeneities; we measure 2.5 fold higher chromatin rigidity near nuclear and nucleolar peripheries than in the nuclear interior. Additionally, we develop an automated light patterning protocol for high efficiency locus repositioning, adapt VECTOR to generate force through a variety of condensate identities, and reposition both chromatin loci and non-chromatin nuclear bodies in living nuclei. Together, these studies build a more complete understanding of internal mechanical processes in the nucleus, and provide a powerful toolkit for the study of cellular organization and function.

## Results

### VECTOR: a rapid and precise system for chromatin locus repositioning in living nuclei

To engineer condensate capillarity, we leveraged our earlier work demonstrating light-dependent condensate induction in the cell nucleus^24,35,39^. We induced proteinaceous condensates in cultured human UOS cells with the two-component Corelet system^39^, comprised of an iLID-GFP-Ferritin 24-mer ‘core’, and an sspB-tagged phase separation-prone IDR. Upon 488 nm light exposure, iLID and sspB interact, decorating each core with up to 24 sticky IDRs, triggering intracellular phase separation. By fusing the same phase separation-prone IDR (e.g. FUS_N_) to a tethering protein that binds to a particular chromatin locus, in this case the TRF1 protein that binds to the repetitive telomeric TTAGGG sequence^40^, we promoted interactions between the condensate and chromatin locus (Figure 1A, B).

**Figure 1.**
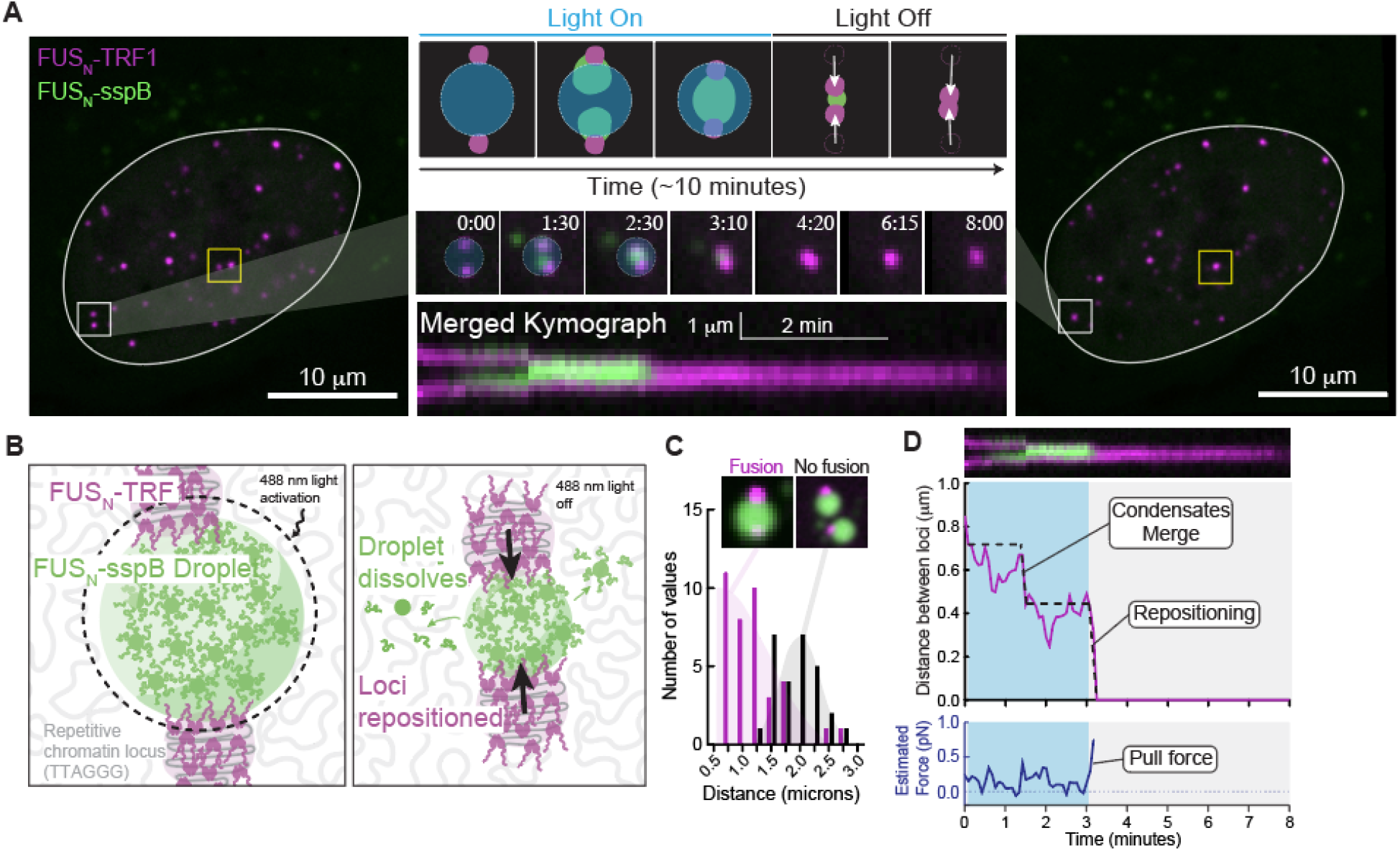
A light-inducible system for rapid, precise repositioning of chromatin loci in living cells. **A**. In a live U2OS cell, two pairs of FUS_N_-miRFP670-TRF1 marked telomeres were targeted for repositioning, boxed in white and yellow (see also Figure S1A). Light activation (blue circle) triggers FUS_N_-sspB synthetic condensate formation at targeted loci. Stills from the white boxed area over time show Corelet condensates (FUS_N_-sspB, green) growing at the locally activated loci (1:30), merging (2:30), repositioning the attached loci as the condensate shrinks (4:20) and the loci remaining in their new positions for the duration of the movie (8:00). A kymograph demonstrates this process over time. **B**. Schematic of VECTOR constructs: TRF1 binds to repetitive telomeric TTAGGG sequences, and the covalently attached FUS_N_ IDR interacts with FUS_N_-sspB within the light-induced Corelet condensate, creating interfacial adhesion. When the light is de-activated (right), the Corelet condensate dissolves but FUS_N_-TRF1 stays attached to the shrinking condensate, bringing together the two associated chromatin loci. **C**. A histogram of distances between pairs of telomeres colored by whether the locus-associated condensates were able to fuse (magenta) or not (black). **D**. Graph of the distance between two targeted loci (top) and estimated pull force applied to the loci (bottom) over time. Blue shading indicates light activation.

By activating blue light on a small region within the nucleus (1.2 micron diameter circle), we can readily nucleate condensates at chosen loci. With maintained light activation, the chromatin-tethered condensates grow and coalesce into one larger condensate, which remains associated with the target loci. Upon light de-activation, the condensate shrinks, leading to a pulling force that repositions the loci. In this example cell, we were able to perform two simultaneous repositionings (white box in Figure 1A, and yellow box in Figure S1A). With this localized light activation protocol, we achieved loci-spanning condensates with diameters up to three microns, enabling successful repositioning of telomeres across multiple microns of nuclear space. However, the probability of condensate merging reduces significantly if loci are separated by more than two microns (Figure 1C), a limitation set by achievable size of the loci-spanning condensate, which could be tuned through changing construct expression levels.

Tracking telomeres during the light activation/de-activation sequence reveals their movement toward one another. We observe a jump together when the two loci-associated condensates merge (0.72 microns/min), then the loci travel towards each other as the condensate shrinks (1.18 microns/min). Directed locus repositioning occurs over 1-2 minutes as the synthetic condensate dissolves, and the loci maintain their new positions for multiple minutes after the synthetic condensate is dissipated, indicating successful long-term repositioning (Figure 1D). Notably, these micron-scale, rapid, and directed movements are achieved entirely without use of ATP-driven motors–the most commonly described source of intracellular force generation. Instead, the forces underlying movement of these genomic loci are generated through surface tension, also known as capillary forces.

The scale of capillary forces is set by the relevant interfacial tension(s), which represent an energetic cost per unit area, with forces generated as a consequence of the geometric drive to minimize surface area^32^. Measuring interfacial tension is challenging, particularly for biomolecular condensates, with most measurements to date made on simplified in vitro condensate systems^7,41,42^. To determine the magnitude of the capillary forces generated in cells with VECTOR, we thus focused on calculating them using a microrheological approach, building from a framework recently deployed to analyze intracellular force generation by magnetic tweezers^31^. In particular, we estimated the force applied on these loci during the light activation/de-activation sequence by calibrating a diffusive Rouse polymer model on tracked telomere mean squared displacements (MSDs), then calculating the forces required to move loci on their observed trajectories^31^. This approach yields estimates of VECTOR forces on the order of ∼0.5 pN; interestingly, this is comparable to the scale of forces generated by individual molecular motors (e.g. kinesin, dynein, and myosin)^43^. This force calibration allows us to monitor the development of these forces over time, during both condensate coalescence and dissolution (Figure 1D, see Supplemental Information).

**Figure S1.**
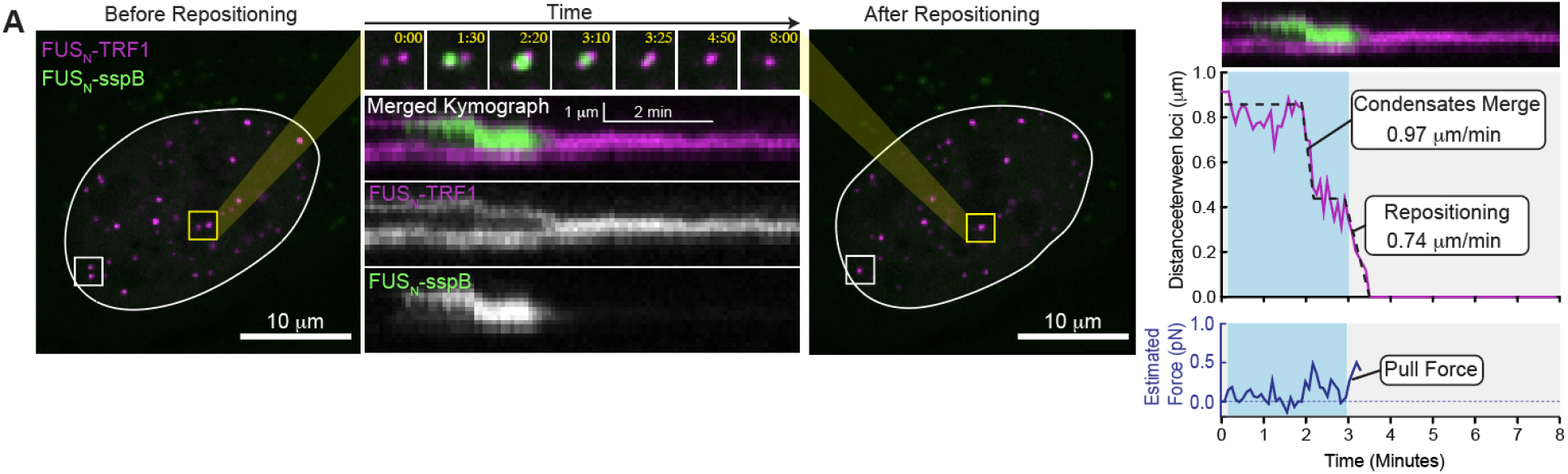
A programmable system for repositioning genomic loci; Related to Figure 1. **A**. Similar to Figure 1: Stills, kymographs and locus distance over time graph for a second locus repositioning (yellow box) in the same cell, demonstrating that multiple pairs of loci can be manipulated at once.

### Adhesion between condensate and chromatin is essential for precise force application

We designed VECTOR such that the capillary forces generated by the shrinking FUS_N_ condensate are transmitted to the target chromatin locus by adhesion between the condensate and the FUS_N_-coated telomere surface. To confirm that such adhesion is necessary, we performed the same light activation/de-activation protocol in cells expressing miRFP670-TRF1, which binds to telomeres but is not tethered to the FUS_N_ IDR (Figure 2A,B). As expected, in this case no sustained force is exerted on the loci, and they are not pulled in together by the shrinking condensate (Figure 2C).

**Figure 2.**
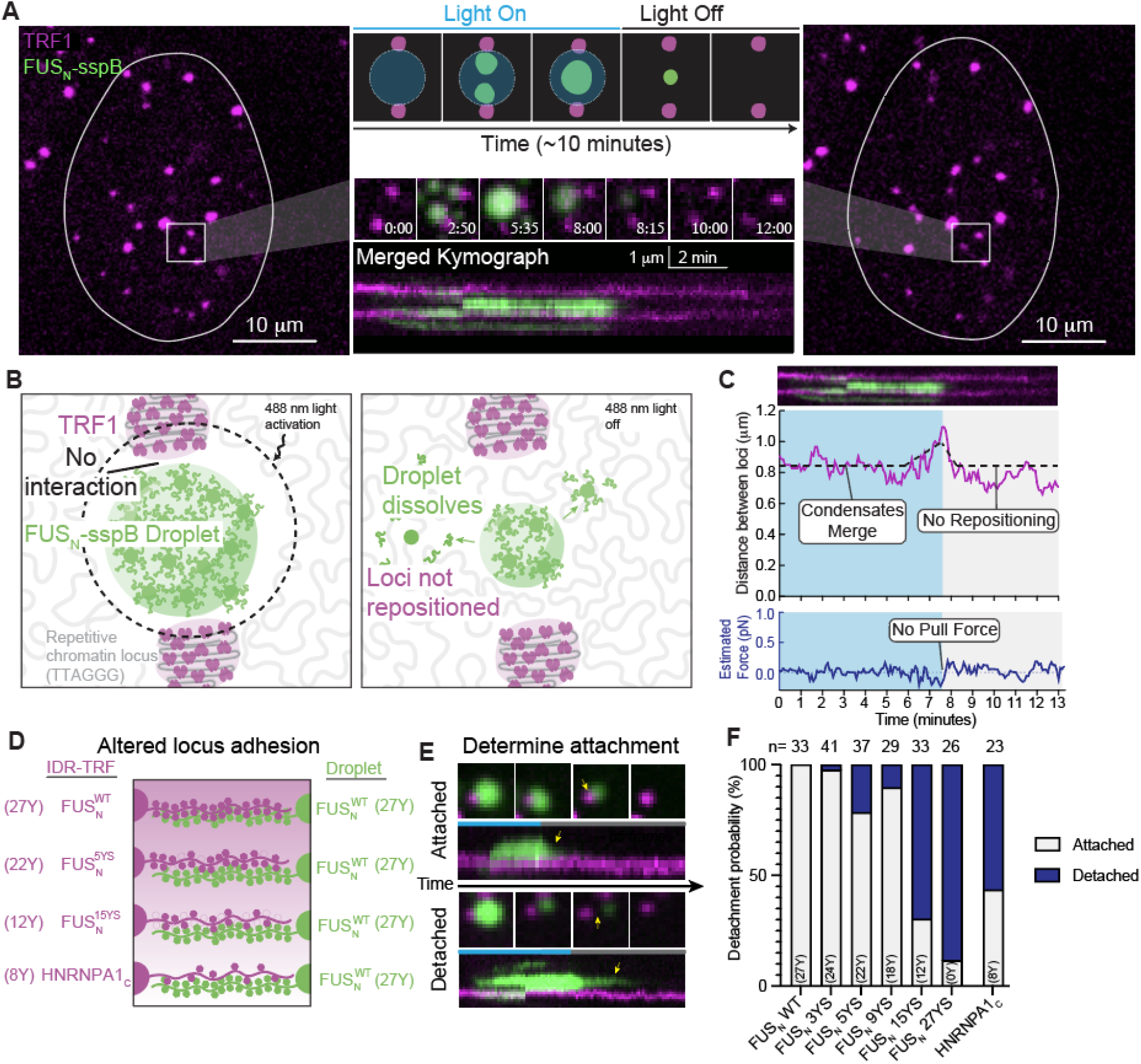
Chromatin-condensate adhesion is required for chromatin locus repositioning. **A**. Still images and kymograph of a nucleus expressing TRF1 without the FUS_N_ IDR, which generates no interfacial interactions between the Corelet condensate and the chromatin locus. The same light activation/de-activation pattern creates condensates that grow (2:50) and fuse (5:35) but do not reposition the chromatin loci when shrinking (8:15-12:00). Kymograph of the two loci (magenta) remaining separate throughout this experiment. **B**. Schematic of the VECTOR constructs with no adhesion between the chromatin locus and Corelet condensate. TRF1 binds telomeric repeat TTAGGG, but does not have any added IDR for interaction with the light-induced Corelet condensate, and so is not repositioned when the condensate is dissolved. **C**. Graph of the distance between two telomeres without condensate adhesion. **D**. Schematic of altered adhesion constructs of telomere-bound FUS_N_ IDR with tyrosine-to-serine (YS) mutations. **E**. Locus-attached condensates were de-activated and classified as staying ‘attached’ if they maintained contact with the locus (top) or as ‘detached’ if they lost contact (bottom). **F**. Quantification of the number of ‘attached’ and ‘detached’ chromatin-condensate pairs with increasing Y-to-S mutations. Trend is statistically significant by two-tailed Pearson correlation, R^2^ = 0.8833, p = 0.0053.

The small expansion and contraction of the distance between chromatin loci without chromatin-condensate adhesion (Figure 2C) may be due to the condensate itself taking up space between the loci. To examine this possibility, we marked all chromatin using SiR-DNA, and locally activated FUS_N_ Corelet condensates. Consistent with previous studies^24,25^, bulk chromatin is excluded from the FUS_N_ condensate as it grows (Figure S2A). Measuring the distance between two endogenous chromatin-poor features (yellow arrows, Figure S2A) over time shows that they are moved slightly apart during activation and recover to their original position after condensate dissolution (Figure S2B). These data suggest that the movement of chromatin loci without adhesion is due to bulk material displacement, and that precise repositioning of the loci is dependent on interfacial interaction with the FUS_N_ condensate.

The amino acids that contribute to self-interaction underlying phase transitions of FUS_N_ and related IDRs have been extensively studied^44-45^, revealing that the 27 tyrosines of FUS_N_^WT^ act as cohesive ‘sticker residues’ separated by ‘spacer residues’ (Figure 2D). To examine the role of tyrosines in facilitating chromatin-condensate adhesion in VECTOR, we created FUS_N_-miRFP670-TRF1 constructs with 3, 5, 9, 15 or 27 tyrosine-to-serine mutations (Y-to-S, Figure 2D-F) and asked whether these mutations led to lowered chromatin-condensate adhesion by their probability of detachment. We observed single locus-condensate pairs during condensate dissolution and determined them to be either ‘attached’ or ‘detached,’ based on whether the centroid of the telomere and that of the condensate moved toward each other or not upon light de-activation (Figure 2E).

We find that detachment of FUS_N_^WT^-tagged telomeres from FUS_N_^WT^ condensates is extremely rare, with 0 out of 33 loci detached (i.e. 0% detachment rate, Fig 2F). Detachment becomes more likely with increasing Y-to-S mutations, resulting in 88% detachment rate of the fully mutated FUS_N_^27YS^-miRFP670-TRF1 construct (Figure 2F); this suggests tyrosines mediate almost all of the FUS_N_^WT^ adhesion energy, which arise from an estimated several hundred IDR-IDR interactions at each chromatin locus (see Supplemental Information). To test whether the number of tyrosines is deterministic of the adhesion strength for a distinct pair of sequences, we utilized telomere-tethered HNRNPA1_C_, an orthogonal IDR that contains 8 tyrosines and has been reported to interact weakly with FUS_N_^40,46^. If the number of tyrosines in a telomere-tethered IDR is the sole determinant of adhesion energy, we would expect HNRNPA1_C_-tagged telomeres to have a detachment rate of about 80%. However, we find only 13 out of 23 (56%) detach, which suggests that spacing of tyrosines, neighboring amino acid context, or other sequence features are important for dictating intermolecular interaction strength, in accord with previous reports^47^.

**Figure S2.**
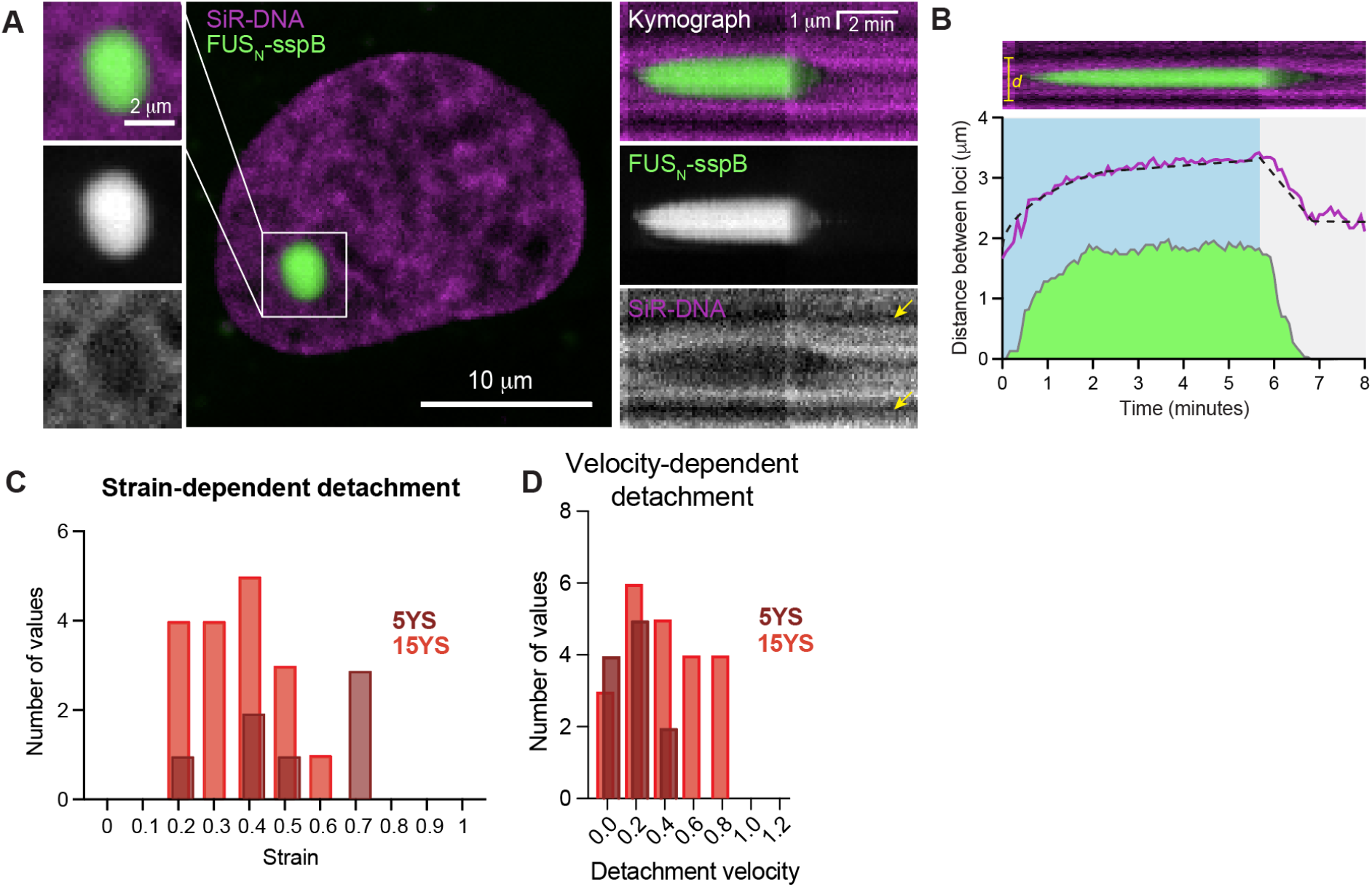
Chromatin-condensate adhesion is required for chromatin locus repositioning and detachment events support a viscoelastic liquid model of chromatin material state, related to Figure 2. **A**. Marking all chromatin with SiR-DNA reveals that chromatin (magenta) is excluded from the bulk of IDR-based condensates (FUS_N_-sspB, green) and kymograph shows that chromatin can relax to fill the void as the condensate shrinks. **B**. Plot of distance between two chromatin-poor nuclear features (*d*, yellow distance) shows they are moved apart slightly as a condensate grows between them, and relax back to their original positions upon condensate dissolution (magenta). In green is the diameter of the condensate between the tracked features. **C.-D**. Detachment probability plotted as a function of strain (**C**) and locus movement velocity (**D**) for chromatin loci attached to FUS_N_^WT^ condensates by FUS_N_^5YS^- and FUS_N_^15YS^-miRFP670-TRF1 (n = 28, 31 nuclei for FUS_N_^5YS^, FUS_N_^15YS^). Strain trends are not statistically significant by two-tailed Pearson Correlation, p = 0.9150, 0.1129 for 5YS, 15YS. Velocity trends are not statistically significant by two-tailed Pearson Correlation, p = 0.0753, 0.6574 for 5YS, 15YS.

### Chromatin is a viscoelastic liquid with local heterogeneities

Since condensate detachment from the target chromatin loci is mediated by stresses generated through deformation of the associated chromatin network, we reasoned that these detachment events might yield insight into chromatin material properties. In particular, if chromatin behaves as a purely elastic material, we would expect the condensate to detach from the locus if it moved far from its initial starting position, when the strain is highest. However, we did not observe a significant positive correlation between strain and detachment probability (Figure S2C). This suggests that chromatin is not a purely elastic material, but rather may exhibit some significant fluid-like dissipation. Materials with partially liquid-like behavior are characterized by stresses which depend not only on the magnitude of deformation, but also on the rate of change of deformation (e.g. velocity or strain rate). With VECTOR, the moving locus follows the receding condensate surface during deactivation, therefore velocity is dependent on the rate of condensate diameter shrinking, which we observed to range between 0.1 and 1.2 microns/minute (Figure S2D). Chromatin loci attached to condensates with full adhesion (FUS_N_^WT^-miRFP670-TRF1) did not detach from the shrinking condensate at any tested velocity (Figure S2D), while loci attached by the very-weak-adhesion construct FUS_N_^15YS^-miRFP670-TRF1 showed a high probability of detachment at all tested velocities (Figure S2D). Loci attached with the moderately mutated construct FUS_N_^5YS^-miRFP670-TRF1 exhibited an apparent velocity-dependent detachment, with detachments more frequent at lower velocities (< 0.5 microns/min); however, we note that the trend was not statically significant (p = 0.0753), and interpretation of velocity dependence is confounded by the fact that the pulling velocity is coupled to the size-dependent condensate dissolution rate. Nonetheless, these data point to chromatin exhibiting significant liquid-like properties, rather than being a purely elastic material.

The relative importance of elastic vs liquid properties is also expected to manifest in whether and how detached loci spring back towards their original positions. To examine this within a theoretical framework, we first built an analytical simulation which mimics dissolution-dependent repositioning by light-inducible condensates (green) wetting a second phase (magenta, representing the chromatin locus) (Figure 3B, see Supplemental Information). In this initial simulation, the condensate forms via uphill diffusion upon light-induced association of the core and IDR, with the free energy of mixing defined according to the Flory-Huggins theory^48^. As we observed in living cells with VECTOR, these simulations show that both condensate coalescence and dissolution lead to the associated loci moving toward each other (Figure S3A), with adhesion required for locus repositioning (Figure S3B). Additional characterization and parameters of these simulations is described in the Supplemental Information (Figures S3C-G).

**Figure 3.**
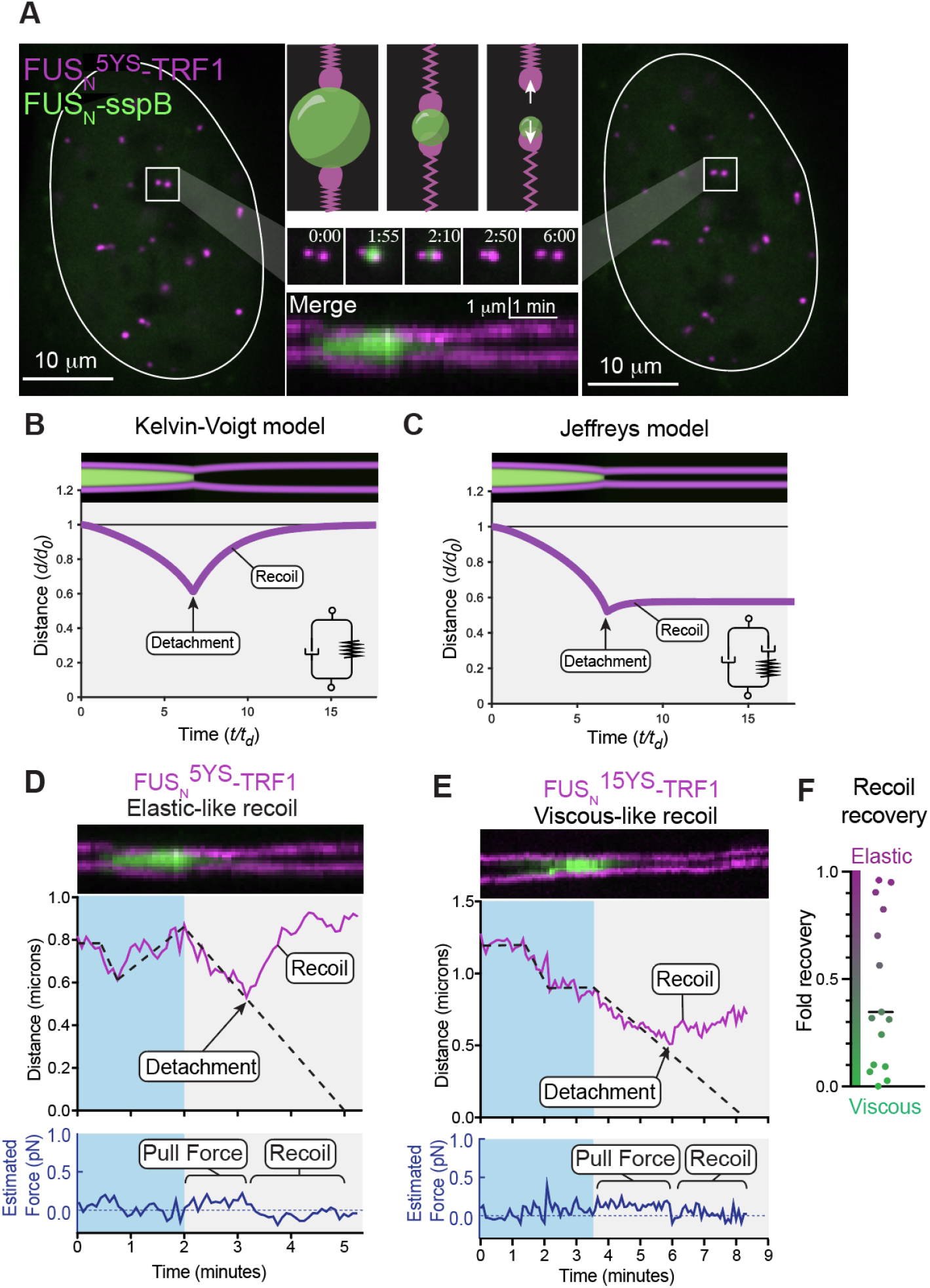
An analytical simulation predicts viscoelastic mechanical properties of chromatin repositioning. **A**. VECTOR experiments using detachment-prone FUS_N_-miRFP670-TRF1 mutants result in detachment and recoil of the associated locus. **B**. Simulating locus detachment from a shrinking condensate and subsequent recoil with a viscoelastic solid (Kelvin-Voigt) model predicts that, during recoil, the chromatin loci always return to their original positions, as the energy of the elastic spring must be dissipated. **C**. Simulating locus detachment and recoil in a linear fluid Jeffreys model predicts that the chromatin loci may not return all the way to their original positions, as some of the energy can be dissipated in the viscous dashpot. Graphs of locus distance and estimated force over time from experiments in live cells show that some detachments result in elastic-like recoil (**D**) while others result in viscous-like recoil (**E**). **F**. Graph of locus’s recovered distance normalized to their original position (fold recovery) for 15 detachments shows a wide spread of both viscous-like and elastic-like recovery behavior.

We then added linear viscoelasticity to the simulation, which gives rise to resistance to locus movement and also impacts the nature of locus detachment and recoil during repositioning (Figure 3B-C). Starting with a system of two loci attached to a single condensate, we simulated the chromatin locus movement during the force of repositioning and considered the chromatin material to be either (i) a viscoelastic solid following the Kelvin-Voigt model, which effectively anchors the locus at its original position by a spring and dashpot in parallel (Figure 3B, graph inset), or (ii) a viscoelastic liquid following the Jeffreys model^49^, which effectively adds a dashpot in series with the spring (Figure 3C, graph inset). While not the only possible models of viscoelastic solids and liquids, we chose these as our initial models because they differ only by an added dashpot.

The Kelvin-Voigt and Jeffreys models make two very different predictions about whether and how loci recoil after they detach from the condensate. In the Kelvin-Voigt model, the single spring spanning the unit dictates that the loci will always return to their original positions (Figure 3B). By contrast, in the Jeffreys model, a detached locus would not necessarily return all the way back to its original position (Figure 3C), since some of the energy in the spring is dissipated by the dashpot; the shorter the spring relaxation time compared to the repositioning time, the less recoil of the locus. We observed locus detachments with the FUS_N_^5YS^- and FUS_N_^15YS^-miRFP670-TRF1 weak-adhesion mutants (Figure 3A), and tracked recoil behaviors. Interestingly, we observed both elastic-like recoil, with detached loci returning entirely to their original positions (Figure 3D), as well as viscous-like recoil, with detached loci returning only partially to their original positions (Figure 3E). Over 15 detachment events using the FUS_N_^15YS^-miRFP670-TRF1 mutant, we observed a wide spread of recoil recovery, with some loci presenting more viscous-like recoveries (green, Figure 3H bottom) and others presenting more elastic-like recoveries (magenta, Figure 3H top). Taken together, our data are most consistent with the Jeffreys liquid model, since it can give rise to varying degrees of recoil, depending on the relaxation timescale. However, these data also indicate significant heterogeneity in the viscoelastic response of the chromatin network.

**Figure S3.**
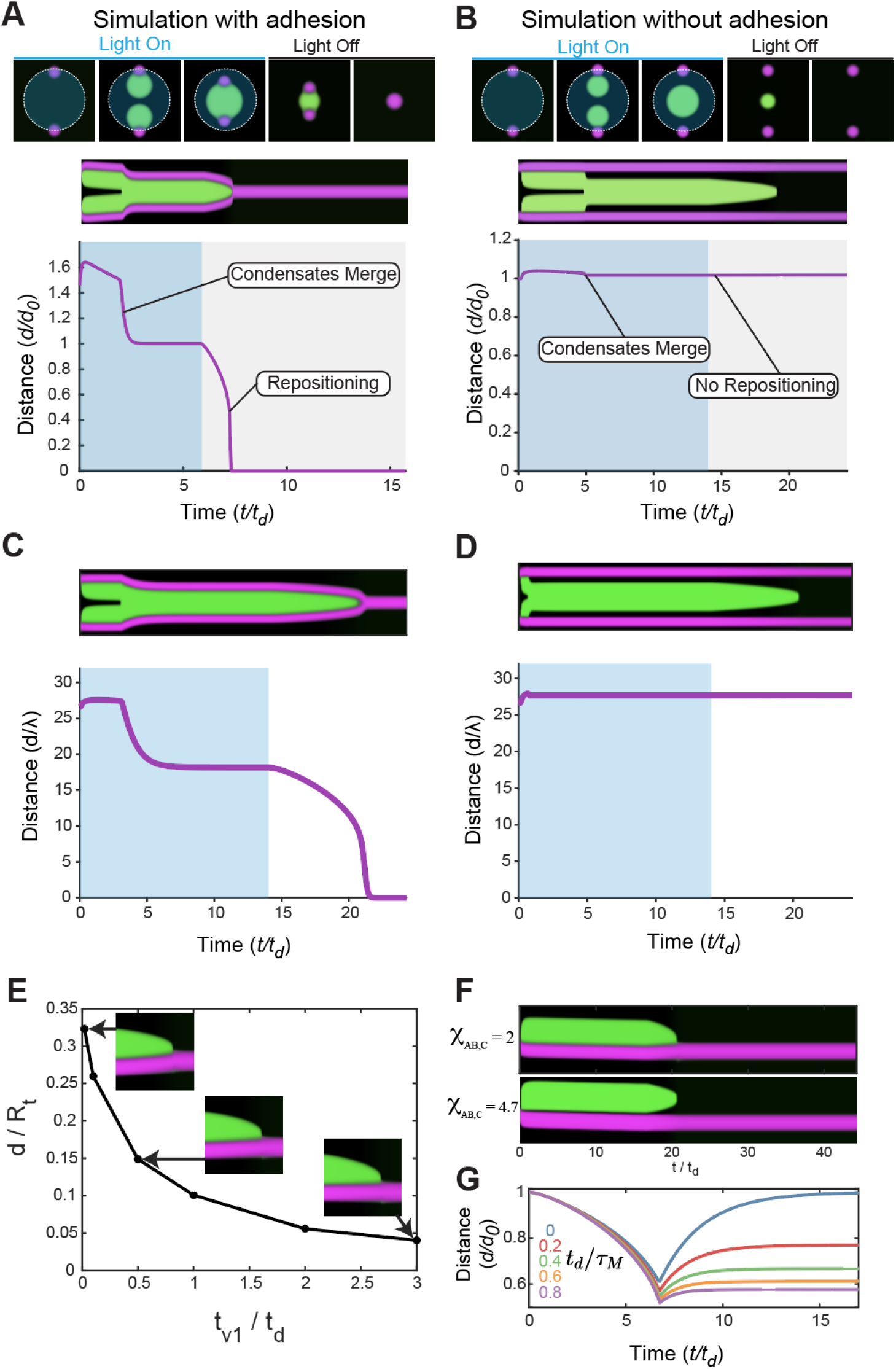
Simulations mimic adhesion-dependent loci repositioning by light-inducible condensates, related to Figure 3. **A**. Still images, kymograph and distance over time plot from simulations of growing and dissolving a central condensate (green) in a two-phase system with adhesion are able to recreate condensate-based repositioning of the second phase (magenta). **B**. Without adhesion, the simulated second phase is not repositioned, in line with experiments. **C-D**. The kymograph and distance between the centers of the telomere loci using identical parameters except for (**C**) *χ*_*AB,C*=_2 and (**D**) *χ*_*AB,C*=_3.5. **E**. Length of telomere displacement normalized by initial telomere radius *R*_*t*_ as a function of *t*_*v1*_*/t*_*d*_. **F**. Kymograph of Corelet nucleation and dissolution when it is next to a single telomere, while *χ*_*AB,C*=_2 (top) and *χ*_*AB,C*=_4.7(bottom). **G**. The distance between two telomeres (normalized by their distance at the moment the light is turned off) over time as the Corelet condensate dissolves at various *t*_*d*_/τ_*M*_. When telomeres detach from the condensate, they recoil.

### Proximity to nuclear or nucleolar periphery dictates chromatin viscoelasticity

To examine the potential origins of chromatin heterogeneity, we tracked diffusion of telomere loci in U2OS cells every 20 seconds for 30 minutes and plotted their pairwise MSDs (Figure S4A). Consistent with previous reports^25,35,50^, we find that, on average, telomeres exhibit constrained diffusion, with an average diffusive exponent, α ≈ 0.6. Since the nuclear periphery is known to be enriched in heterochromatin, which is more densely compacted and thus presumably stiffer, we plotted these pairwise MSDs binned into two categories by their nuclear location (Figure S4A); peripheral (blue), or internal (mauve). Peripheral pairs exhibit lower MSD magnitude than internal pairs, consistent with local heterogeneity in chromatin mechanics across the nuclear interior, and higher viscoelastic resistance near the nuclear periphery (Figure S4B).

Given these indications of heterochromatin-associated mechanical heterogeneity, we reasoned that upon pulling on two loci associated with different local mechanical environments (e.g. peripheral heterochromatin vs. internal euchromatin), we may be able to use VECTOR to directly observe this heterogeneity across small (e.g. 1-2 μm) length scales. To examine the feasibility of this concept, we again used our simulations to predict the movement of chromatin loci, either in a mechanically symmetric or asymmetric system (Figure 4A). Here, we define a simplified dimensionless parameter, ρ, which represents the mechanical resistance of the spring-and-dashpot viscoelastic system, with ρ_1_ andρ_2_ indicating resistance of the two respective loci attached to a singular condensate. In the symmetric case ρ_2_ /ρ_1_ =1, the viscoelastic parameters between the two loci are equal, which result in symmetric locus movement, each traversing 50% of the distance between them. By contrast, when we simulate the movement of loci in an asymmetric system with unequal viscoelastic resistance (ρ_2_ /ρ_1_ =5), we find an unequal locus movement (77%, 22%; Figure 4A, bottom and Figure 4B). A series of simulations (ρ_2_ /ρ_1_ =2,4,6,8,10) reveal the linear relationship between differential viscoelastic properties and the ratio of distance traveled by each locus (*d*_*2*_*/d*_*1*_) (Figure 4C). Further analysis of the relationship between viscoelastic heterogeneity and asymmetric locus movement is provided in the Supplemental Information (Figures S4D-K).

**Figure 4.**
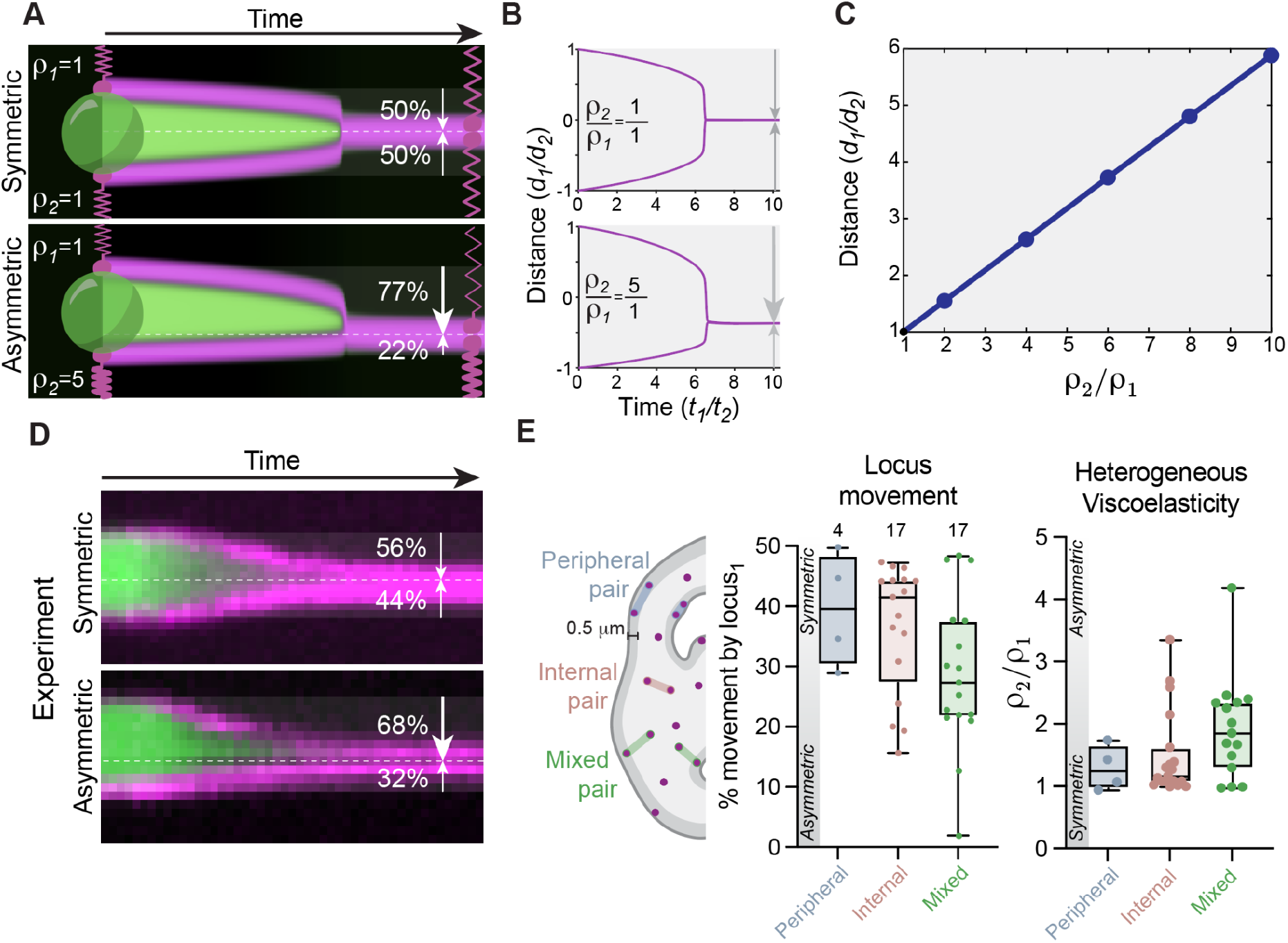
Proximity to nuclear or nucleolar periphery dictates local heterogeneity in chromatin viscoelasticity. **A**. Simulations of shrinking condensates with equivalent viscoelastic resistance of the two attached loci (ρ2/ρ1=1) result in equal distance traveled by each locus (top; 50%, 50%), while simulations with unequal viscoelastic resistance (ρ2/ρ1=5) show unequal distance traveled by each locus (bottom; 77%, 22%). **B**. Plots of the loci positions over time in simulations of a symmetric system (top) or asymmetric system (bottom). **C**. Graph describing the relationship between the ratio of viscoelastic strength of two loci (ρ2/ρ1) and the ratio of the distance traveled by each locus (*d*_*1*_*/d*_*2*_). **D**. Experiment-derived kymographs of a symmetric VECTOR system (top) and an asymmetric VECTOR system (bottom). **E**. Quantification of asymmetric locus movement in pairs of telomeres both < 0.5 µm from the periphery of nucleus or nucleolus (Peripheral, blue, n = 4), both internal (mauve, n = 17), or one peripheral and one internal (Mixed, green, n = 17). Statistical significance was calculated using one-way ANOVA with multiple comparisons, ns = not significant. Locus movement: Peripheral-Internal p = 0.8906. Peripheral-Mixed p = 0.2248. Internal-Mixed p = 0.1304.

Consistent with local heterogeneity in mechanical response manifesting in unequal loci displacement, we frequently observe both symmetric (Figure 4D, top) and asymmetric (Figure 4D, bottom) movement of chromatin loci pairs. Moreover, we find that when both loci are nuclear ‘internal’ (> 0.5 microns from a nuclear or nucleolar periphery) or both loci are ‘peripheral’ (< 0.5 microns from a nuclear or nuclear periphery), movement is relatively symmetric, while ‘mixed’ pairs (one peripheral locus and one internal locus) tend to be more asymmetric (Figure 4E). Given the linear relationship between differential viscoelastic properties and the ratio of distance traveled by each locus established in simulations (Figure 4C), these data imply that ‘mixed’ pairs connect loci from environments with greater viscoelastic heterogeneity (ρ _2_/ρ_1_ ≠1 ; Figure 4E). Indeed, in 16 out of 17 ‘mixed’ pairs of VECTOR loci tested, the nuclear-internal locus (locus_2_) moves > 50% of the distance, consistent with lower viscoelastic resistance than its peripheral partner locus (locus_1_) (Supplemental Figure 4C). Together, these results support a model of a viscoelastic liquid nuclear interior with up to 2.5 fold increased resistance within 0.5 microns of nuclear and nucleolar peripheries.

**Figure S4:**
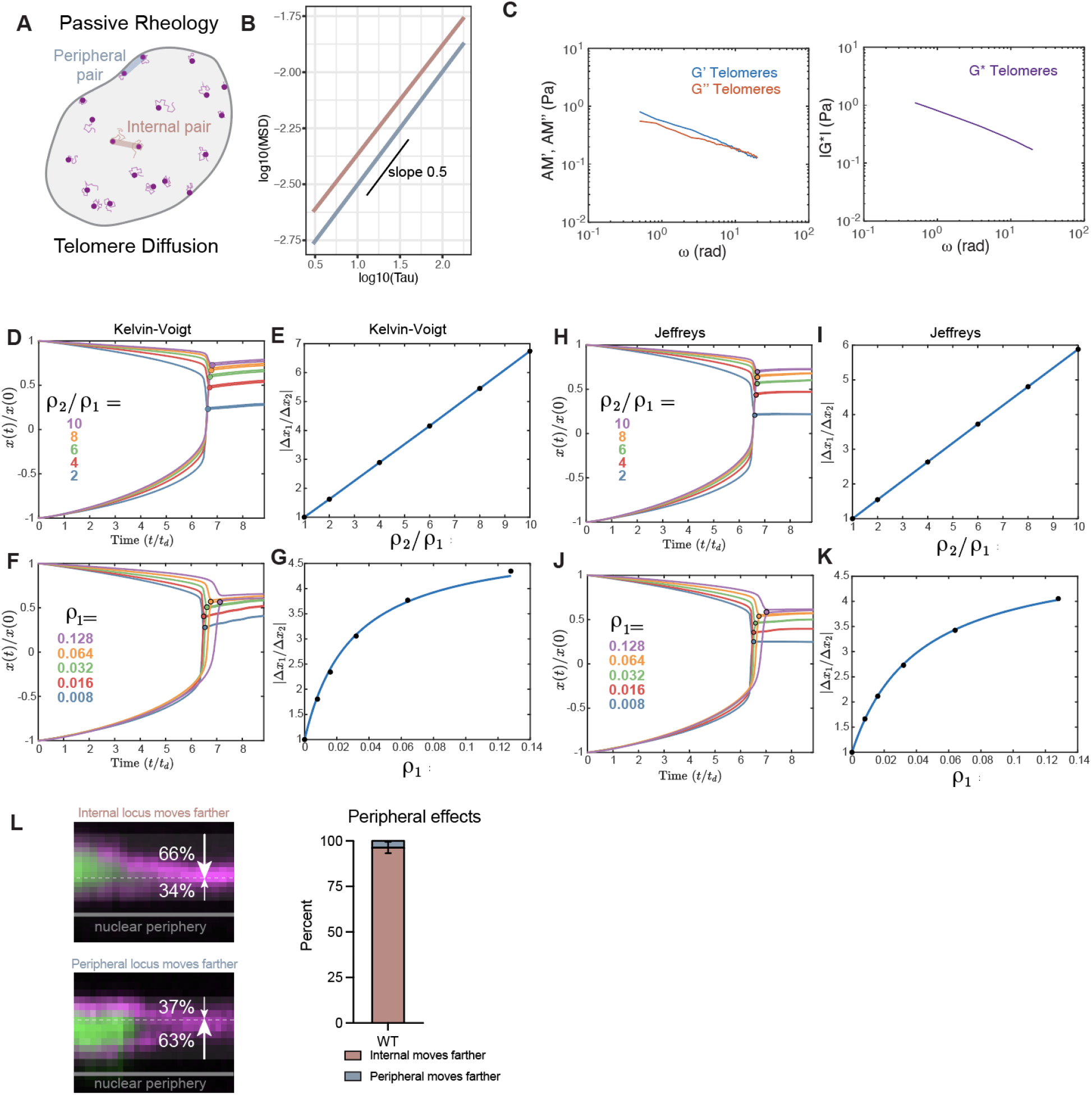
Rheological measurements and simulations of chromatin material state, related to Figure 4. **A**. Schematic of definitions of Peripheral (blue) and Internal (mauve) pairs of telomeres for MSD analysis. **B**. MSD curves of Internal, Mixed, or Peripheral pairs of diffusing telomeres. Log_10_ timestep in seconds, n = 11 cells. **C**. Apparent Moduli (AM) of elastic (AM’) and viscous (AM’’) components of chromatin viscoelasticity calculated from MSD traces (left). Viscoelastic modulus G* obtained from apparent moduli (right). Using the Kelvin-Voigt model with τ_1_ /*t*_*d*=_τ_2_/*t*_*d*=_2, (**D**) the trajectories of the two telomeres *x*(*t*)during the dissolution of the Corelet droplet and (**E**) the ratio of displacement of the two telomeres as a function of ρ_2_/ρ_1_ with fixed ρ_1 =_0.05. The displacement is defined to be the distance between the telomere position at t = 0 and its position at the moment when the two telomeres merge, indicated by the filled circles in (**D**). The values of the displacement ratio |Δ*x*_1_ /Δ*x*_2_| as shown as black dots in (**E**). Also using the Kelvin-Voigt model, (**F**) and (**G**) are the trajectories and ratio of displacement as a function of ρ_1_ with fixed ρ_2_/ρ_1 =_5. Using the Jeffreys model with τ_1_ /*t*_*d*=_τ_2_/*t*_*d*=_2 and τ_*M*,1_ /*t*_*d*=_τ_*M*,2_/*t*_*d*=_2, (**H**) and (**I**) are the trajectories and ratio of displacement as a function of ρ_2_/ρ_1_ with fixed ρ_1 =_0.05, (**J**) and (**K**) as a function of ρ_1_ with fixed ρ_2_ /ρ_1_ =5. The blue lines and curves in (**E, G, I, K**) are fitted to the black data points using Eq. S21 (Supplemental Information). **L**. Example kymographs with nuclear periphery marked (gray) showing an example where the nuclear-internal locus moves toward the peripheral locus (top) and an example where the peripheral locus moves toward the nuclear-internal locus (bottom). Quantification of the probability that in a mixed pair, the internal or peripheral locus moves farther (N = 3 experiments, n = 6, 6, 5 nuclei per replicate).

### Versatility and Programmability of VECTOR

The data shown above all rely on cell-specific, user-defined laser activation and de-activation at particular genomic loci of interest, an approach limited in throughput capacity. To scale up the number of successful repositioning events while maintaining the specificity and precision required for identifying, activating, and moving loci, we tested a series of automated protocols that incorporate real-time feedback into region-of-interest (ROI) generation for light patterning (see Methods and also Supplemental Information; Figure 5A). Global activation creates temporally controllable but small condensates that mostly fail to merge. A thinner activation region within the nucleus induces larger condensates, and sliding this activation box across the nucleus can promote condensate fusion, but this approach is relatively slow (t > 60 min per frame), due to the numerous sequential activation/de-activation cycles required per field of view. An array pattern of punctate activation regions (i.e. square lattice) creates large condensates that occasionally merge, but are not all associated with target loci.

**Figure 5.**
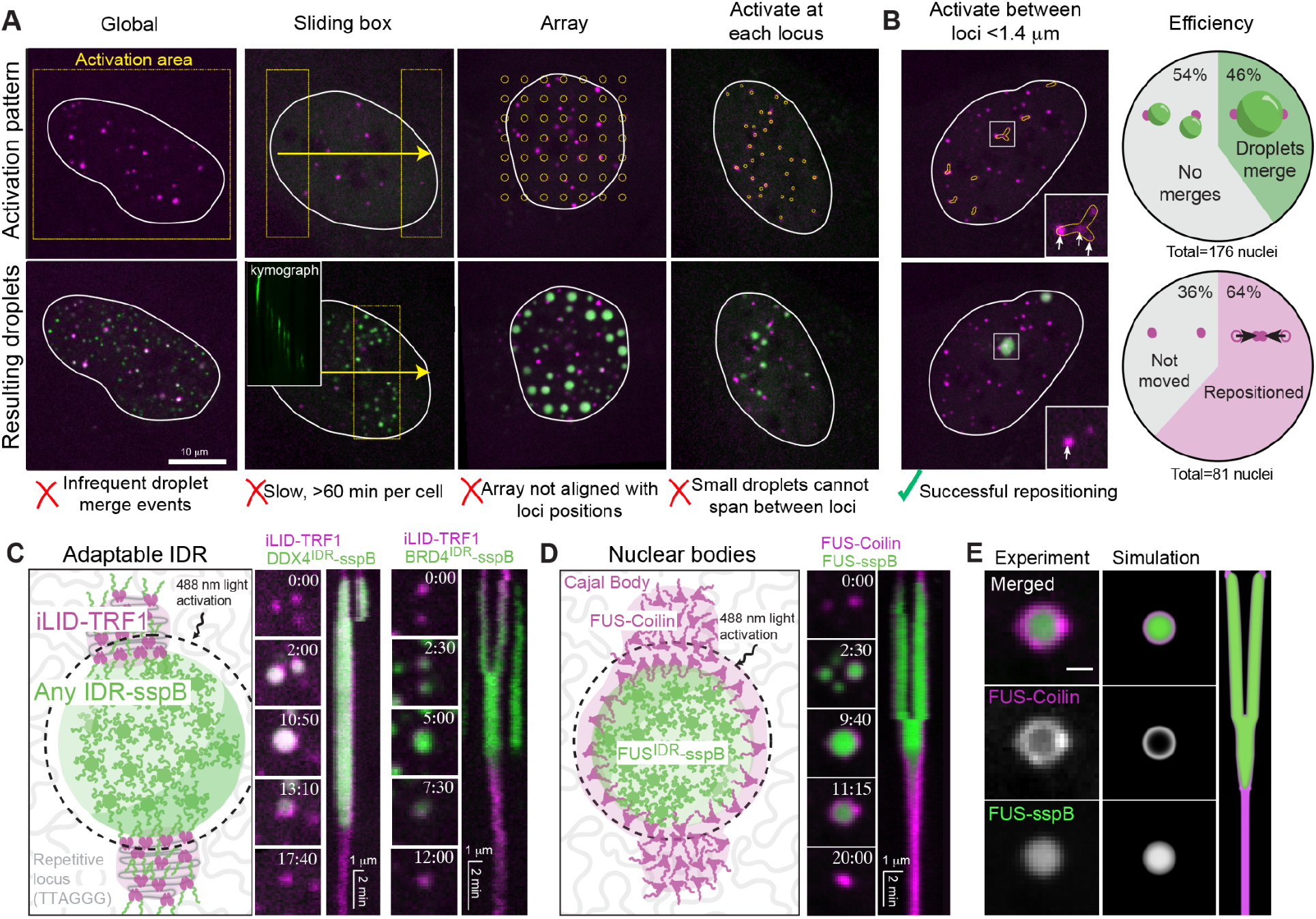
Versatility and Programmability of VECTOR. **A**. Images with automatically-generated activation regions indicated in yellow (top) and resulting condensate pattern (bottom) for light patterns: Global, Sliding Box, Array patterning, and Activation at each locus. **B**. Left: example image of automatically-generated activation regions between loci <1.4 microns apart using a custom JOBS protocol (see Supplemental Information). RIght: Quantification of automated light-patterning protocol efficiency. Of 176 nuclei attempted, 81 nuclei or 46% produced productive condensate merger events, and of those 52 nuclei or 64% resulted in successful locus repositioning. **C**. Schematic of VECTOR with adaptable IDR capability; iLID-miRFP670-TRF1 (magenta) recruits Any-IDR-mCherry-sspB (green) upon light activation, creating adhesion between the repetitive locus and IDR-based condensate. Example stills and kymographs of successful locus repositioning using iLID-TRF1 with DDX4^IDR^-mCherry-sspB or BRD4^IDR^-mCh-sspB. **D**. Schematic of VECTOR system that repositions non-chromatin nuclear bodies: Cajal bodies. FUS_N_-miRFP670-Coilin (magenta) marks Cajal bodies and creates adhesion between them and a FUS_N_ Corelet condensate (green). Two FUS_N_-miRFP670-Coilin bodies (magenta) are repositioned. Merged kymograph shows fusion of the two Cajal bodies. After repositioning, Cajal bodies remain fused until the end of the observation period (20:00). **E**. Stills of individual channels of FUS_N_-miRFP670-Coilin (magenta) and FUS_N_ condensate (green) from both experiment and simulation (including kymograph, right), exhibiting their core-shell architecture.

Having determined the limitations of these different activation protocols, we find that the most successful approach is using real-time image analysis feedback to identify close telomere pairs, then automatically create activation regions at the identified loci, leading to directed condensate formation at target loci. These automatically activated locus-associated condensates successfully coalesced between loci at most 1.4 microns apart, similar to the probability limit of condensate coalescence we observed with our manual targeting strategy (i.e. Figure 1C). Therefore, we added a step to the automated protocol that calculates the distance between all locus pairs, and creates activation regions connecting loci at most 1.4 microns apart (Figure 5B). Bright spot detection followed by activation between nearby loci is consistent and efficient, resulting in at least one pair of loci associated with the same condensate in 46% of cells (81 out of 176 nuclei attempted) and leading to scalability and successful repositioning in 64% of those nuclei (52 out of 81 potential locus pairs) in less than five minutes per field of view (Figure 5B).

Given the ability of this feedback protocol to program VECTOR for high throughput applications, we next sought to examine VECTOR’s generalizability with respect to the details of condensate/locus targeting. We first sought to manipulate the position of not just telomeres but any target genomic sequence, using an engineered dCas9-based chromatin-condensate adhesion construct. Building from the previously described CasDrop system^24^, we utilized an enzymatically dead Cas9 to bind chromatin and covalently attached an iLID to accumulate FUS_N_ IDRs upon blue light activation and create chromatin-condensate adhesion. We targeted dCas9-HaloTag-iLID to telomeres with a guide RNA that recognizes TTAGGG repeats, visualized telomeres with Halo-JF647, and applied the localized blue light activation/de-activation sequence (Figure S5A). With this dCas9-based system, we find the chromatin-tethered loci quickly detach from condensates, precluding sustained force generation. This may suggest that the current dCas9-based approach provides insufficient adhesion between synthetic condensates and the target genomic locus, a problem that should be addressable with further construct engineering to increase IDR density or adhesion strength.

Until this point, we have utilized a specific FUS_N_ IDR to mediate interfacial force generation. However, capillary forces should be a general feature of condensates, not dependent on any single protein. To confirm that the repositioning force of VECTOR is not dependent on the FUS_N_ IDR specifically, we replaced the chromatin-tethered FUS_N_ IDR with iLID to create iLID-miRFP670-TRF1^35^ (Figure 5C). This construct binds to telomeric TTAGGG sequences, recruits IDR-mCh-sspB during the light activation sequence, and can be paired with non-FUS_N_ IDR constructs including DDX4^IDR^-mCh-sspB and BRD4^IDR^-mCh-sspB (Figure 5C, S5B). Similar to the FUS_N_ system, during activation, the chromatin loci remain adhered to the condensate surface, and upon de-activation the iLID-miRFP670-TRF1 construct leads to successful locus repositioning (Figure 5C, S5B).

Finally, we attempted to reposition non-chromatin nuclear objects, focusing on Cajal bodies. We exchanged the TRF1 telomere-binding protein with a Cajal body protein, Coilin, to create FUS_N_-miRFP670-Coilin that mediates adhesion between a Cajal body and FUS_N_ Corelet condensates (Figure 5D). Interestingly, when we applied the localized light activation/de-activation pattern, FUS_N_-miRFP670-Coilin distributes to wet the entire surface of the FUS_N_ condensate. Nevertheless, from two original Cajal bodies, only one is left when de-activation is complete, indicating successful repositioning. In parallel, we adapted our simulation to mimic interfacial interaction of the Corelet condensate with a nuclear body-associated protein rather than chromatin-bound protein by allowing for mobility of the simulated FUS_N_-miRFP670-Coilin, which yielded its redistribution over the surface of the simulated Corelet condensate, recapitulating the core-shell architecture we observed in cells (Figure 5D, S5C-D). These data demonstrate that VECTOR is capable of repositioning nuclear bodies in addition to chromatin loci, providing a facile way of rapidly re-programming nuclear body organization.

**Figure S5:**
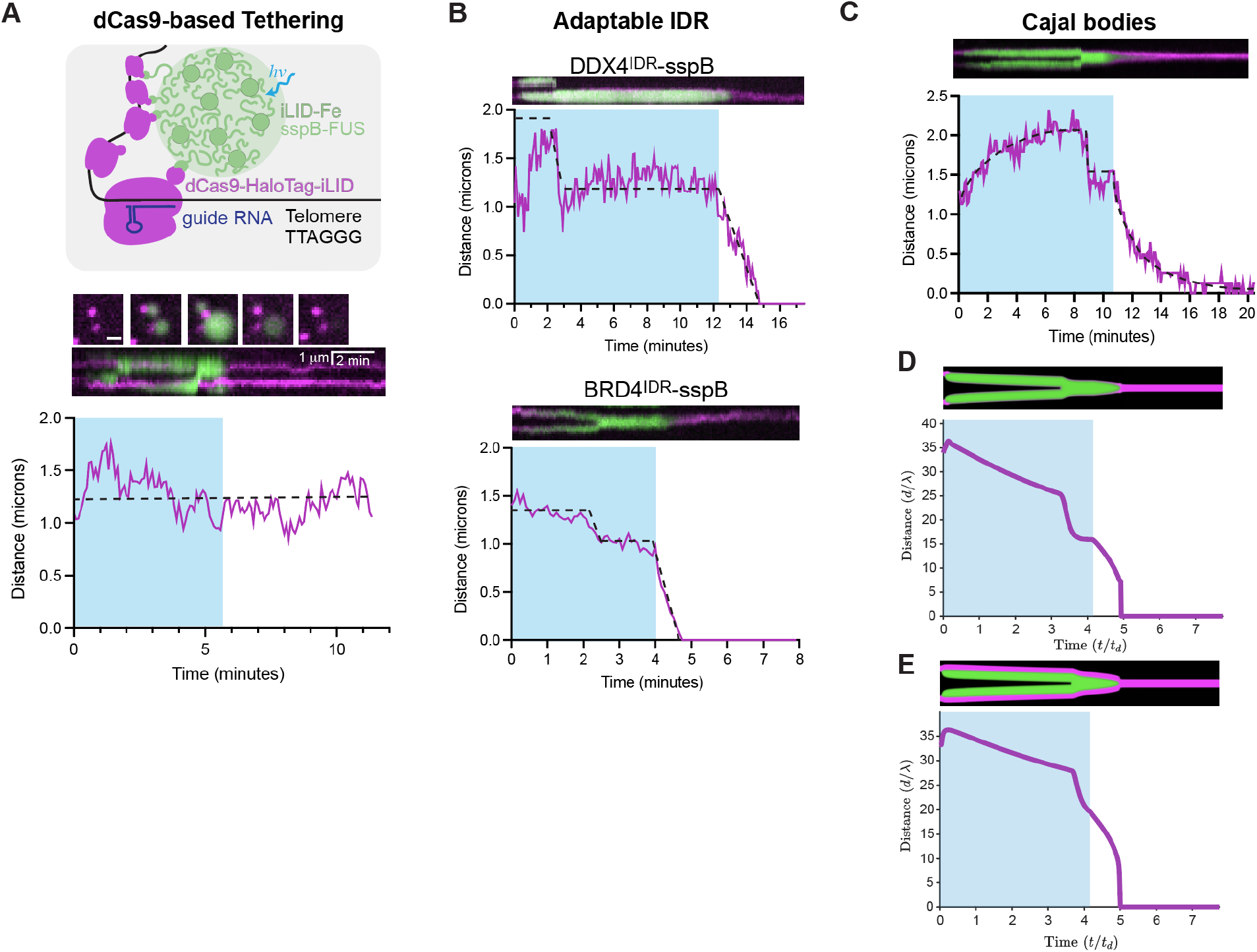
Versatility and programmability of VECTOR; related to Figure 5. A. Schematic of dCas9-based tethering between chromatin and condensate; an sgRNA recognizing telomeric TTAGGG guides dCas9-HaloTag-iLID to bind telomeres, and was visualized with Halo-JF647. Upon blue light (hv), FUS_N_-sspB binds both to dCas9-tethered iLID and Ferritin core iLID to create chromatin-tethered condensates. Perhaps due to lack of sufficient adhesion, light de-activation leads to condensate dissolution but no force generation. **B**. Distance graphs for adaptable IDRs DDX4 and BRD4; related to Figure 5C. **C**. Kymograph and distance between Cajal bodies over time in experiment (**C**) and simulation (**D**). To simulate Cajal bodies, simulation in (**D**) has no interaction kernel, so the ‘coilin’ (magenta) is distributed over the surface of Corelet condensate (green). **E**. Simulated kymograph and distance between magenta loci using identical parameters as (**D**) except adding back interaction kernel.

## Discussion

In this work we developed VECTOR, a technique that deploys synthetic light-controlled condensates to impart force on specific chromatin loci in living cells and consequently reposition them while sensing the local microenvironment. Using this system, we showed how capillary forces associated with biomimetic condensates are capable of generating pN-level forces in living cells and harnessed these physical forces to characterize distinct material responses across different nuclear positions. Our findings on (1) lack of significant correlation between degree of displacement and detachment probability, (2) velocity rate-dependence of detachment probability, and (3) high frequency of incomplete locus recoil, altogether suggest that the material state of chromatin in live nuclei is consistent with the viscoelastic fluid-like Jeffreys model, exhibiting a high degree of heterogeneity in viscoelastic resistance, which is coupled to nuclear- and nucleolar-proximity (Figure 6). Our findings underscore the complexity and heterogeneity of chromatin material state within the nucleus, which likely arise from a variety of sources, including differences in compaction, epigenetic modifications and/or protein binding.

**Figure 6.**
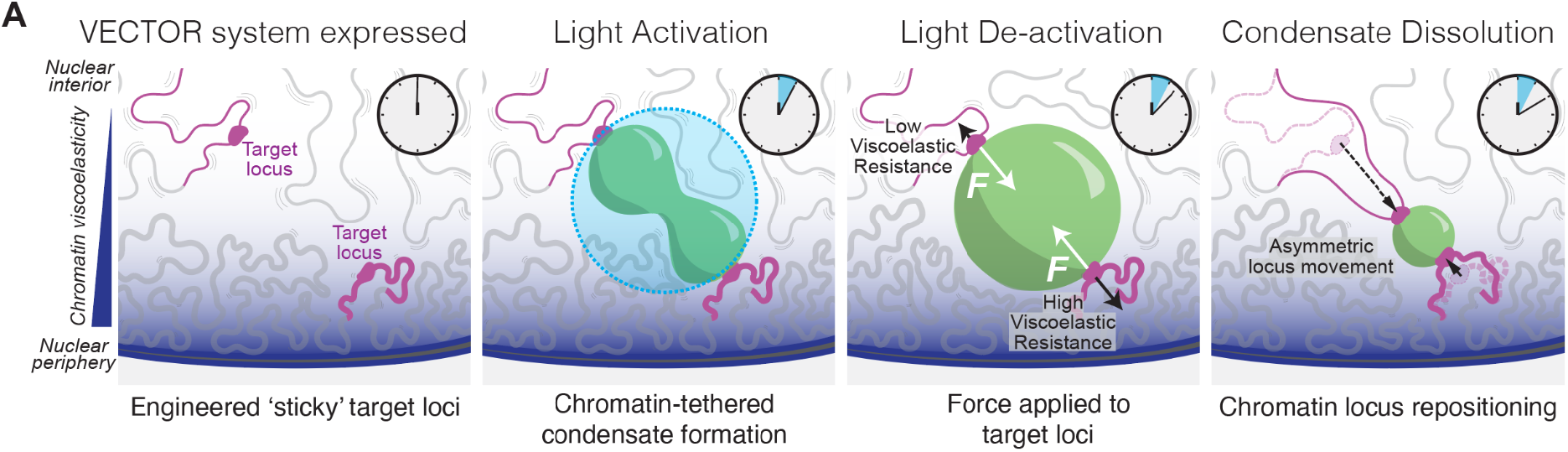
Interfacial interactions mediate precise force generation in living cells. **A**. Target chromatin loci are bound by VECTOR system proteins, creating seeding sites for light-inducible condensates to grow and fuse during a few minutes of localized blue light activation. Upon light de-activation, target loci remain attached to the shrinking condensate due to interfacial interactions, which leads to equal forces applied on the attached loci. Uneven viscoelastic resistance of the attached loci leads to asymmetric repositioning events, revealing increased local chromatin viscoelasticity near the nuclear periphery.

With VECTOR, we were able to reposition telomere loci across microns of space in just seconds-to-minutes, using condensate-associated interfacial forces. Our data indicate that these forces can be appreciable, with 0.1-1 pN-level forces that are comparable to those generated by individual molecular motors (e.g. kinesin or dynein). We were able to harness and direct interfacial forces via IDR-mediated adhesion between the condensate and targeted telomeres, which were ideal loci because their repetitive nature makes them efficient nucleation seeds for synthetic condensates^24,40^, and their abundance in the cell makes it highly likely to find two in a suitable position for synthetic condensate coalescence. In addition to repositioning telomeres, we were also able to physically deform other nuclear structures, such as Cajal bodies (Figure 5D), suggesting that a variety of cellular structures can be sites of intracellular force generation.

A desirable future application of VECTOR is the ability to reposition any locus of interest through dCas9-based genomic targeting strategies (Figure S5A). However, our initial experiments in this direction resulted in insufficient chromatin-condensate adhesion, perhaps due to a lower density of dCas9 than TRF1 molecules binding per sequence length, leading to fewer IDR-IDR interactions at the chromatin-condensate interface. Ultimately, the VECTOR system requires not only targeting of chromatin-condensate interaction, but also a sufficient degree of adhesion during force application; how interfacial forces manifest in such adhesivity represents a largely unexplored aspect of condensate biology, with broad potential implications in intracellular organization. We estimate that the interfacial interactions required are on the scale of a few hundred FUS_N_ IDR-IDR interactions (see Supplemental Information); when this adhesion is compromised by mutating a significant percentage of the tyrosine “stickers,” the condensates typically detach prematurely (Figure 2D-F). Future studies will adapt the dCas9-based VECTOR constructs used here to apply force on any locus of interest, beyond telomeres, and potentially incorporate alternate strategies to increase condensate-locus adhesivity. These adaptations to VECTOR will be key for further studies examining the causality of pairwise locus interactions, locus positioning relative to nuclear bodies/compartments^51,52^, and dynamics of functional outcomes.

Our demonstration of condensate-mediated genome reorganization utilizes interfacial forces at the pN-level, a magnitude comparable to ATP-fueled molecular motors, but in this case does not require ATP^32^. Instead, energy that underlies these forces is stored in the condensate interface, which before coalescence and rounding represents a non-equilibrium thermodynamic state, and controlled through blue light activation of the iLID-sspB interaction^39^. Upon removal of light, genomic loci repositioning follows the surface of the shrinking condensate, and therefore is dependent on the rate of the condensate diameter shrinking, which we find to be generally between 0.75 and 1.25 µm/min. This speed is comparable to whole-chromosome movement in mitosis, but not as rapid as individual molecular motors such as kinesin, which walk along microtubules at 2-3 µm/sec^53^.

Notably, our data clearly show that work can be generated by dissolving condensates, reminiscent of the finding that disassembling microtubules also generates significant force^54^. In both cases, sustained attachment to the object against which force is applied is key (e.g. Dam1 in the case of MTs^54^, IDR-mediated adhesion in the case of our VECTOR system), highlighting the importance of interfacial interactions in force application. In the context of the nucleus, this suggests that both rapid assembly and disassembly of endogenous, chromatin-bound condensates may represent an additional layer of spatiotemporal regulation of pairwise locus interactions and genome organization.

Chromatin loci experiencing forces via VECTOR show force-response and recoil behavior consistent with a viscoelastic liquid Jeffreys model of chromatin material state. In previous studies, the discrepancy between lack of intermixing of chromatin with itself (e.g. labeled histones or replication-labeled DNA) and rapid FRAP recovery of chromatin-binding proteins (e.g. heterochromatic factors) have been interpreted as a solid-like chromatin scaffold surrounded by a nucleoplasmic liquid^28-29^. However, in this work, we have applied pN-level point forces to individual chromatin loci and directly measured the mechanical response, illustrating that at the mesoscale (0.1-1 micron), chromatin material state has both viscous and elastic components. In particular, we find that after cessation of force application, loci do not necessarily return to their original positions, indicating a partially dissipative viscoelastic liquid rather than a pure elastic solid model of chromatin material state. A viscoelastic liquid model is consistent with historical data, including constrained and coordinated diffusion of chromatin^55-56^ and scale-dependent protein mobility within the chromatin environment^57-58^.

Importantly, we measured mechanical heterogeneity within the nucleus up to 2.5-fold more rigid near nuclear and nucleolar peripheries than in the interior, which is consistent with known subnuclear localization of heterochromatin^59^, signifying that chromatin’s epigenetic state may significantly influence local viscoelasticity. This is consistent with increased chromatin compaction of heterochromatic sequences^60^, which may underlie local heterogeneities. Increased rigidity at the nuclear periphery compared to interior would explain observations of high mechanical stiffness of the whole nucleus^61-62^, even while interior genomic elements may be associated with significant chromatin rearrangement, for example upon transcriptional activation.

This work represents the first deployment of synthetic condensates for the systematic and programmable control of intracellular forces, and their use in interrogating mechanical properties of the genome. We have described the utility of interfacial adhesion between condensates and cellular objects to do organizational work within living cells, and suggest that endogenous condensates may utilize their interfacial interactions for similar purposes. Our results highlight the ubiquitous nature of such intracellular interfacial forces, and their potential importance in regulating structural changes of the 3D genome, and likely a variety of other intracellular structures. The interplay between forces generated by condensates, and the mechanical resistance of cellular structures, provides an exciting new perspective on the rich regulatory landscape underlying chromatin compartmentalization, nuclear organization, and their consequences for cellular physiology and disease.

## Acknowledgements

We thank Leonid Mirny, Antoine Coulon, Simon Grosse-Holz, and Veer Keizer for helpful discussions regarding estimating pull force applied on chromatin loci and for sharing their ‘Rousepull’ code (https://github.com/OpenTrajectoryAnalysis/rousepull), Mikko Haataja for helpful advice on the modeling, Shunsuke Shimobayashi for the FUS_N_-miRFP670-TRF1 and HNRNPA1_C_-miRFP670-TRF1 constructs, Mackenzie Walls for the FUS_N_ mutant original vectors, David Sanders for the FM5 vectors, Jing Xia for the array patterning protocol, Lennard Wiesner for thoughtful comments on the manuscript, Evangelos Gatzogiannis for microscopy assistance, and other members of the Brangwynne Lab for helpful feedback and discussions.

This work was supported by the Howard Hughes Medical Institute, the Princeton Biomolecular Condensate Program, the Princeton Center for Complex Materials, a MRSEC (NSF DMR-2011750), and the AFOSR MURI (FA9550-20-1-0241). A.R.S. is a Life Science Research Foundation fellow through the Mark Foundation for Cancer Research (No. AWD1006303). Y.K. is supported by the NSF GRFP (DGE-2039656). H.Z. is supported by the Princeton Bioengineering Initiative Innovators (PBI^2^) Postdoctoral Fellowship.

## Contributions

A.R.S., Y.K., and C.P.B. designed the study, with A.R.S. and Y.K. performing experiments and analysis, with advice from C.S. and H.Z. H.Z. created experiment-parallel simulations, with advice from A.K. N.O. performed telomere MSD tracking and pairwise data analysis. Y.-C.C. developed original technologies that provided the basis for these experiments. A.R.S., Y.K., and C.P.B. wrote the manuscript, and A.R.S. and Y.K. made the figures, with contributions from all authors.

## Methods Cell Culture

All cell lines were incubated in and grown at 37°C with 5% CO_2_. U2OS cells that were obtained from the ATCC (authenticated via ATCC’s STR profiling) were cultured in DMEM (GIBCO, 11995065) with 10% FBS (Atlanta Biological, S11150H) and 1% streptomycin and penicillin (GIBCO, 15140122), grown at 37°C with 5% CO_2_.

### Construct design and cloning

FUS_N_-miRFP670-TRF1 and HNRNPA1_C_-miRFP670-TRF1 constructs^40^ were kind gifts from Shunsuke Shimobayashi. FUS_N_ point mutants were PCR-amplified from original vectors^39^ gifted by Mackenzie Walls, using CloneAmp HiFi PCR Premix (Takara Bio 639298). These PCR fragments were inserted into linearized FM5 lentiviral vectors that carry standardized linkers that were kind gifts from David Sanders^63^, using the In-Fusion HD cloning kit (Takara Bio, 638910).

In addition to these point mutants, all other DNA fragments of interest were cloned using the same protocol and reagents. All constructs were confirmed by GENEWIZ Sanger sequencing.

### Constructs

FUS_N_-miRFP670-TRF1^40^

FUS_N_^3YS^-miRFP670-TRF1

FUS_N_^5YS^-miRFP670-TRF1

FUS_N_ ^9YS^-miRFP670-TRF1

FUS_N_ ^15YS^-miRFP670-TRF1

FUS_N_ ^27YS^-miRFP670-TRF1

HNRNPA1_C_-miRFP670-TRF1^40^

iLID-miRFP670-TRF1^35^

FUS_N_-miRFP670-Coilin

DDX4-mCherry-sspB

BRD4^ΔN^-mCherry-sspB

### Lentivirus production and lentiviral transduction

All live cell experiments used cells stably transduced with lentivirus. Lentiviruses were produced by plating Lenti-X 293T cells (Takara Bio, 632180) into 6-well plates to reach ∼70% confluence at the time of transfection. 24-48 hours after plating, the transfer plasmid and helper plasmids VSVG and PSP were transfected into the Lenti-X cells using Transit293 transfection reagent (Mirus, MIR 2700) incubated in OptiMEM (modified from ^63^). ∼48 hours after transfection, viruses were harvested and filtered using a 0.45 μm filter (Pall Life Sciences) and were either then used immediately or stored at -80°C. U2OS cells plated at 30-50% confluency in 96-well glass-bottom plates (Cellvis) were transduced for 2-3 days before live-cell imaging experiments.

### Microscopy

Cells for all live cell imaging experiments were plated on 96-well glass-bottom plates and incubated at 37°C and 5% CO_2_ by an Okolab microscope stage incubator with a 96-well insert. Images were taken on a spinning disk (Yokogawa CSU-X1) confocal microscope with an Andor DU-897 EMCCD camera on a Nikon Eclipse Ti body and a 100x oil immersion Apo TIRF objective (NA 1.49 MRD01991), and a Nikon LU-NV laser launch. A second spinning disk confocal microscope with a Nikon Plan Apo VC 100x 1.4 oil immersion objective, Nikon LU-N4 laser launch, and Oko Labs Bold Line Cage Incubator with a 96-well plate insert with 0.1% accuracy CO_2_ control at 37°C was used to take optogenetic stimulation images and the automated imaging experiments. 488 nm, 561 nm, and 640 nm lasers were used to image mGFP, mCherry, and miRFP670 constructs, respectively, on both microscopes.

### Optogenetic stimulation

Specific regions in the nuclei of cells were locally activated by using a Mightex Polygon digital micromirror device (DMD) to pattern blue light (488 nm) activation from a Lumencor SpectraX light engine. U2OS cells expressing FUS_N_-miRFP670-TRF1, NLS-GFP-iLID-Fe and FUS_N_-mCherry-sspB were imaged using a specific local activation protocol: 1) Pre-activation, imaging the mCherry (541 nm) and miRFP670 (640 nm) channels every 5 s for 10 s; 2) Activation, using a circular region of interest (ROI) (with diameter of 1.2 μm) to locally activate two genomic loci/nuclear bodies to seed, grow, and fuse FUS_N_ Corelet condensates using the 485 nm DMD every 5 s for 2-10 min; 3) De-activation, back to only imaging the 561 and 640 nm channels every 5 s to allow the FUS_N_ Corelet condensates to dissolve and pull together attached loci/structures.

### Automated imaging protocol

All automated imaging protocols were created by using the JOBS module of the Nikon NIS-Elements software. All protocols included this workflow: 1) Define well selection, 2) Set up the autofocus, 3) Designate the 541 nm and 640 nm lasers as the ‘Capture Definition,’ and 3) Pre-define points with cells expressing all relevant constructs listed above in the Optogenetic Stimulation section. Each of the following automated imaging protocols used the DMD to stimulate at pre-defined ROIs at 395 nm wavelength at 30% intensity unless noted otherwise:

1. **Sliding box** In order to mimic a slow scan across a cell from left to right, we defined a rectangular box (1.2 μm wide, 64 μm tall boxes) as the ROI and stimulated two of these boxes at a time with one box remaining from the previous activation sequence to maintain condensates formed from the previous sequence and the second box to form new condensates in the current loop). For each predefined point, the JOBS protocol took a Z-stack before and after the optogenetic activation/de-activation segment. After the first Z-stack, cells were imaged with two ROI boxes for 2-5 min (5 sec/frame) using the Capture Definition NDStimulation with Sequential Stimulation feature every second. The ND Acquisition sequence was then used to image cells for the de-activation segment (5 sec/frame) for 5 minutes.
2. **Array patterning** The array patterning protocol used the following parameters on the Polygon pad: 0.405 μm diameter of the stimulation ROI with each ROI 2.835 μm apart from its closest neighboring ROI with the array pattern covering an area of 56.7 μm by 56.7 μm of the field of view. The protocol stimulated the ROIs using the 395 nm wavelength at 100% intensity. Cells were imaged for 10 minutes (5 sec/frame).
3. **All telomere stimulation** All telomeres were detected using the ‘Bright spot detection’ with the NIS-Elements General Analysis 3 program. All bright spots were then converted into ROIs and the same workflow laid out in the ‘Sliding Box’ section was applied.
4. **Detect close telomere pairs** Each telomere was detected similarly as the ‘All telomere stimulation’ protocol. Using General Analysis 3, each bright spot’s centroid x, y positions were measured and two bright spots whose centroids were 10 pixels (1.35 μm) or less apart were connected with a thin line. These measurements made and overlaid a binary image of these connecting lines on the telomere channel that were then converted into ROIs for stimulation.

### Image analysis

#### Kymograph production

Movies of chromatin locus movement were registered to correct for whole cell movement in FIJI (ImageJ 1.52p)^64^ using HyperStackReg^65^, Rigid Body translation. A line was drawn across the activation region and kymographs were created using the MultiKymograph plugin.

### Loci tracking during repositioning

Registered movies of chromatin locus repositioning were cropped to a region containing the relevant telomeres, then loaded into the TrackMate plug-in^66^ in FIJI. Telomeres were identified in the miRFP670 channel with the LoG detector, with an estimated blob diameter of 0.7 microns and no initial thresholding. Spots were filtered by quality to eliminate background. Tracks were generated using Simple LAP tracker with max linking distance 1 micron, gap-closing max distance 1 micron and gap-closing max frame gap 0 frames. Tracks of the relevant telomeres were identified and the XY distance (*d*) between them calculated for each time point 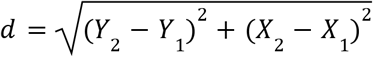 **Detachment probability**

Constructs with FUS_N_ point mutants (FUS_N_^3YS^, FUS_N_^5YS^, FUS_N_^9YS^, FUS_N_^15YS^, FUS_N_^27YS^) fused to miRFP670-TRF1 were created, expressed in living U2OS cells along with Corelet components, and imaged using the Optogenetic Stimulation protocol as described above. ROIs were aimed at singular telomere loci to observe how a single locus-condensate pair would behave during condensate dissolution during the de-activation sequence. Chromatin-condensate pairs were classified as ‘attached’ if the condensate dissolved towards the locus or ‘detached’ if the condensate dissolved concentrically, away from the telomere.

### Detachment probability as a function of strain and velocity

Probability of detachment of single FUS_N_^15YS^-miRFP670-TRF1-marked locus from its attached condensate was separated into bins by strain (0 - 1) and by velocity (0.0 - 0.2, 0.2 - 0.4, 0.4 - 0.6, 0.6 - 0.8 and 0.8 - 1.0 microns/min). Images were processed in FIJI, using Trackmate to identify the XY position of the locus of interest using the miRFP670 channel and XY position of the condensate centroid using the FUS_N_^WT^-mCherry-sspB channel. Strain was defined as (*d*_*0*_*-d*_*d*_)/*d*_*0*_ where *d*_*0*_ is the initial distance between the chromatin locus centroid and condensate centroid during the first frame of deactivation, when force application begins, and *d*_*d*_ is their distance at the moment of detachment, determined visually. Detachment velocity was defined as the linear slope of locus displacement over time for the 10 - 30 seconds before detachment; loci that did not detach were counted as ‘attached’ in each bin within their highest velocity moved.

### Symmetric/asymmetric locus movement characterization

All images were analyzed using FIJI for the image pre-processing steps and Python 3.7.10 for image processing and analysis. During pre-processing, each nucleus was corrected for whole cell movement using HyperStackReg, and set to a contrast level sufficient for segmentation in Python. In Python, the last frame of the activation sequence from the mCherry and miRFP670 channels were used to segment nuclei and telomeres respectively, and repositioned telomeres from the last frame of the de-activation sequence were segmented, then binary nuclear masks were processed with the Canny Edge Detection method to identify the nuclear and nucleolar peripheries. The distance from all edges to telomere centroids in both frames were calculated, and the edge bearing the shortest distance from each telomere was called out and characterized as the point of ‘nearest periphery.’ This information was then used to find the actual distance between the closest edge and telomere of interest 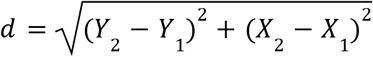. Distances were plotted as a fraction of the total distance between the two telomeres, classified into three distance bins. A telomere pair was classified as ‘peripheral’ if both telomeres were < 0.5 μm from the periphery of the nuclear/nucleolar periphery, ‘internal’ if both were > 0.5 μm from the nearest periphery, or ‘mixed.’

### Heterogeneity in chromatin viscoelasticity

Numerical asymmetric distances moved by each locus were directly converted into a ratio of viscoelastic resistances associated with each locus (ρ_*2*_*/*ρ_*1*_) using the calibration chart provided by a series of simulations (Figure 4C). These were plotted again in categories ‘peripheral’, ‘internal’ and ‘mixed’ as above.

### Mean squared displacement analysis

U2OS cells were transduced with lentivirus to express miRFP670-TRF2, to fluorescently mark telomeres, and imaged 2-3 days post transduction. Telomere tracking movies were obtained by imaging the 670 nm channel every 3 seconds for 30 minutes. MSD data were analyzed as previously described^25^. Briefly, tracking movies were cropped to isolate individual nuclei, and processed in FIJI using Trackmate with blob diameter 0.7 μm and intensity threshold 85. LAP tracker was used to build trajectories with maximum frame-linking and gap-closing set to 0.5 μm. Tracks were exported from Trackmate for further analysis.

Pairwise mean squared displacement (MSD) is defined as a function of time-step τ according to the following equation, where *d(t)* is the distance between the two puncta at time *t*:

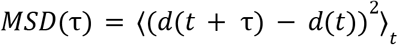

Using a pairwise MSD in lieu of the more traditional single-particle version allows control for translation and rotation of the cell during video acquisition without relying on image registration. A custom script in R^67^ was used to compute and plot MSD (τ) curves. Trajectories shorter than a minimal duration cutoff (1200 seconds) were automatically discarded from the analysis; pairs of loci that coexisted for longer than the minimum cutoff were then identified, and MSD (τ) curves were computed for each pair, with maximum τ set at 80 frames. Pairwise MSDs were binned into ‘peripheral’ and ‘internal’ categories based on their component telomere’s nuclear positioning, determined algorithmically as within 0.45 microns of the nuclear periphery and confirmed visually.

### Generating pull force estimation plots

See Supplemental Information

### Simulations and Modeling

See Supplemental Information

### Statistical analysis

Statistics were performed using GraphPad PRISM version 9.1.0 software (GraphPad). Statistical significance of detachment probabilities in Figure 2 and Supplemental Figure 2 were calculated using Pearson Correlation; R^2^, p value and size of n are noted in figure legends and captions accordingly. Statistical significance of asymmetric locus movement as a function of locus position was calculated using one-way ANOVA with multiple comparisons, number of replicates, and size of n are noted in the figure legends and captions accordingly.

## Supplemental Information

### Passive microrheology to determine viscoelastic moduli

We used telomere mean squared displacement data to estimate the order of magnitude of the nuclear interior viscoelastic modulus, G*, according to Eq. 10-12 in ^S1^.

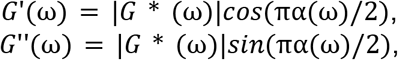

where

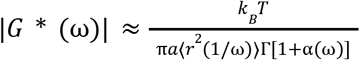

Other studies have used similar approximations on MSD tracking data of peroxisomes from live ATP-depleted cells to estimate the elastic shear modulus G’, viscous shear modulus G”, and viscoelastic modulus G* of the cytoplasm at angular frequencies between 0.126 and 628 rad/s^S2^. We estimated the telomere’s radius to be on the order of 0.1 micron that is consistent with previous studies^S3-S8^, and frame rate was 3 seconds. Using a public MATLAB code (http://measurebiology.org/wiki/MATLAB:_Estimating_viscoelastic_spectrum_using_Mason%27s_method), we calculated from average telomere MSDs of 18 nuclei the apparent elastic and viscous moduli. We call these apparent moduli because the approach assumes thermodynamic equilibrium, so there may be some contribution from out-of-equilibrium (e.g. ATP-dependent) effects. We find the apparent elastic and viscous moduli of chromatin measured via telomere probes are of equivalent magnitudes, with the elastic modulus slightly dominating for lower frequencies. Additionally, we calculate that G* is on the order of 1 Pa (Supplemental Figure 4C) across angular frequencies between 0.355 and 12.58 rad/s.

### Initial order of magnitude force estimation from telomere trajectories

To approximate the magnitude of force applied on a chromatin point locus by a shrinking condensate, we used an estimation method that is based on the generalized Langevin equation under the assumption that thermal fluctuations dominate the noise. From Eq. 1 in^S9^, ignoring inertia and with the presence of the pulling force *F*(*t*), we have

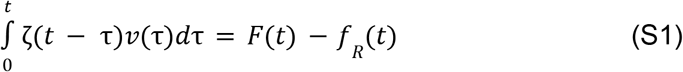

where *t* is time, *ζ*(*t*) is the generalized time-dependent memory function, *v*(*t*) is the particle velocity, *f*_*R*_(*t*) is the random force that satisfies ⟨*f*_*R*_(*t*)⟩ = 0 and ⟨*f*_*R*_(0)*f*_*R*_(*t*)⟩ = *k*_*B*_*Tζ*(*t*), where *k*_*B*_ is the Boltzmann’s constant and *T* is the temperature. Taking the ensemble average of Eq. S1, we have

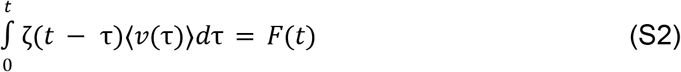

During the pull, the telomere displacement curve in Supplemental Figure 1 shows that the distance between telomere loci approximately decreases linearly in time with a constant velocity *v*_0_ ∼ 1µm*/*min. As an approximation for the ensemble average of the velocity during the pull, ⟨*v*(*t*)⟩ ≈ *v*_0_. Hence, during the pull the force is

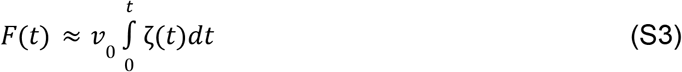

Denoting the Laplace transform with a tilde, the Laplace transform of *F*(*t*) is

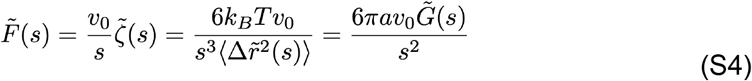

where *s* is the frequency in the Laplace domain, 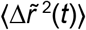 is the mean squared displacement (MSD) of a tracer particle, and 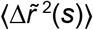 is the corresponding Laplace transform, *a* ∼ 0.1µm is the radius of the telomere, consistent with previous measurements as noted above, 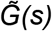 is the Laplace transform of the shear modulus. The last two equalities are from Eqs. 3-4 in ^S9^.

As an order of magnitude estimation, from the MSD measured in Supplemental Figure 4B, at the frequency of 1 min^−1^, which is the frequency scale that corresponds to the time window during which the pull force is applied onto the telomere, we have 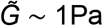 calculated from G* as shown in Supplemental Figure 4C (see ‘Passive microrheology to determine viscoelastic moduli’ section), hence *F* ∼ 1pN, which is on the same order of magnitude as the forces calculated using the Rousepull method from^S10^ (see ’Generating pull force estimation plots’ section).

### Generating pull force estimation plots

In order to estimate the pull force applied on chromatin loci during the dissolution of the condensate, we referred to “Section II. Force Inference” in ^S10^ Supplementary Text and utilized corresponding code for force inference (https://github.com/OpenTrajectoryAnalysis/rousepull) provided by ^S10^ authors. With this “Rousepull” method, we use the Rouse polymer model to infer forces in chromatin loci pulling experiments, calibrating it on our MSD tracking of telomeres in the absence of the condensate (i.e. in the absence of applied force), using prefactor 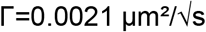 as a first approximation that was extracted from our telomere tracking MSD data 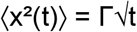 (chromatin loci undergo Rouse diffusion essentially in the absence of the pull force). Using HyperStackReg^S11^ in FIJI, we first corrected for whole cell movement and plotted the relative motion (i.e. distance between two loci) over time (in seconds) to obtain estimated force profiles (Figures 1D, S1A, 2C, 3D-E).

### Physical model

We perform simulations of pulling and merging condensates with the Corelet system via capillary forces, using a phase field model coupled with linear viscoelastic models. The Corelet construct consists of a ‘core’ (A) with 24 binding sites that can bind to the IDR component (B) when light-activated due to iLID-sspB association^S12^. The other component of interest is the telomere and the telomere binding protein (FUS_N_-TRF1), which we treat as a single species that forms a phase that is distinct from the Corelet condensate, and which we intend to move via capillary forces due to its interaction with the condensate. All other species are considered as the buffer (S). We model the mixture using the Flory-Huggins free energy of mixing Δ*g*^S13^. When there is no light activation, A and B do not associate, and thus do not undergo phase separation. When light-activated, A and B bind (A + B → AB), and the majority exists in the associated form AB which forms the Corelet droplet. Hence for simplicity, we set the interaction parameter between A, B, and other species to zero and only consider the interaction parameter between AB, C, and S,

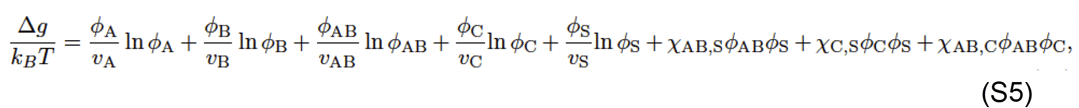

where Δ*g* is the free energy of mixing per lattice site based on the Flory-Huggins lattice theory, *k*_*B*_ is the Boltzmann constant, *T* is the temperature, *ϕ*_*i*_ is the volume fraction of component *i*, where *i* ∈{A, B, AB, C, S, *v*_*i*_ is the number of lattice sites occupied by species *i*, and *χ*_*i,j*_ is the Flory interaction parameter between component *i* and *j*. The volume fractions satisfy the constraint that ^∑^_*i*_ *ϕ*_*i*_ = 1. For a phase-separating system, in addition to the bulk Δ*g*, the total free energy of mixing also includes the contribution from the concentration gradient, which we assume to have the same coefficient *λ*^2^

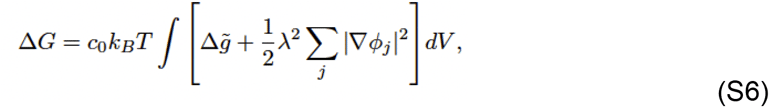

where *c*_0_ is the number density of lattice sites, and 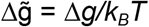 is the non-dimensionalized bulk free energy. This free energy is used by Cahn and Hilliard for non-uniform systems and in Cahn-Hilliard equation^S14^, which is conventionally used to model phase separation. Due to the energy associated with the concentration gradient, the diffuse interface between phases is on the order of *λ*.

We define the chemical potential to be the variational derivative

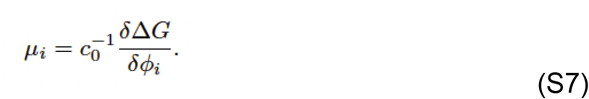

Notice that the variational derivative is taken while satisfying the 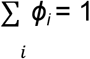 constraint, in other words, the buffer is treated as a reference component whose volume fraction is a function of those of other components 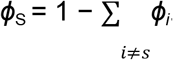. We define the activity *a*_*i*_ by

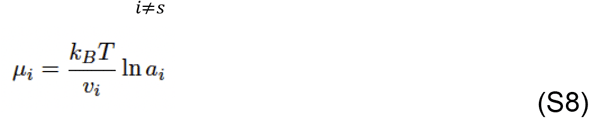

The equilibrium condition for the association reaction A + B → AB is *a*_A_*a*_B_ = *K*_*d*_ *a*_AB_, where *K*_*d*_ is the dissociation constant that changes with the light intensity. The kinetics of association can be described by a rate that follows detailed balance *R* = *k*(*a*_A_*a*_B_ − *K*_*d*_*a*_AB_)^S15^. Both the kinetic prefactor *k* and the dissociation constant *K*_*d*_ depend on the light intensity, which we denote with subscripts “on” and “off” to refer to when the light is on (*R* = *k*_on_(*a*_A_*a*_B_ − *K*_*d*,on_ *a*_AB_)) and off (*R* = *k*_off_ (*a*_A_*a*_B_ − *K*_*d*,off_ *a*_AB_)). Suppose the association is a volume-conserving reaction, that is, *v*_AB_ = *v*_A_ + *v*_B_. As modeled previously for light-activated droplet systems^S16^, the gradient in the chemical potential causes a diffusive flux and we assume a constant mobility *M*_*i*_ for species *i*. In summary, the governing equations for all the species are

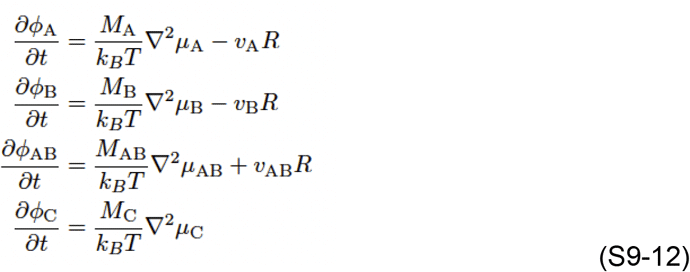

The model above considers C to be a freely moving fluidic species. However, we are also interested in the case where C interacts with certain regions of the chromatin, such as telomeric chromatin. Because TRF1 binds to telomeric DNA, which may experience viscoelastic forces, we model the interaction between *C* and the telomere using an isotropic interaction kernel *K*(|**r** − **r**_*i*_|), which acts as a potential well around the telomere that TRF1 binds to, where **r** is any point in space as defined above, and **r**_*i*_ is the center of the telomere locus *i*. In other words, extending Eq. S6, the total free energy is now

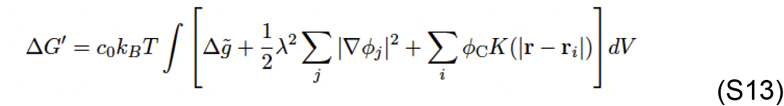

where the summation over *i* refers to all the telomere loci of interest. Here, we use a Gaussian interaction kernel *K*(|**r** − **r**_*i*_|) = *U*_0_ exp−*∥***r** − **r**_*i*_*∥*^2^*/λ*^2^, which has the same length scale as the diffuse interface. Note that *U*_0_ is dimensionless.

Due to the interaction between the telomere and the droplets, the droplets also exert a force **F**_*i*_ on the telomere,

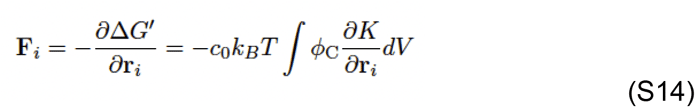

The equation of motion of the center of the telomere locus can be described using various viscoelastic models which we study later:

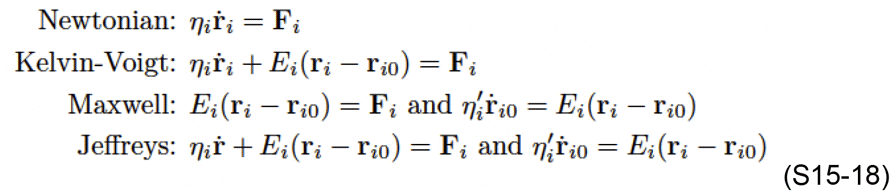

where the dot refers to the time derivative, *E*_*i*_ is the stiffness constant of the spring element, *η*_*i*_ is the friction coefficient (inverse mobility) of the dashpot, 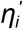 is the friction coefficient of the additional dashpot in series with dashpot in the Maxwell and Jeffreys models.

The time scales of these models are important to consider and have significant impact on the dynamics. Therefore, we define them here. Because we are interested in the dissolution and coalescence time of the Corelet condensate, we define time scales based on *R*_0_, defined to be half the distance between the two telomere loci which are located on opposite sides of and in contact with the synthetic condensate, or approximately the radius of it.

The diffusion-limited dissolution/growth time scale of the condensate can be derived based on the Cahn-Hilliard equation (eliminating the reaction term in Eq. S11) to be 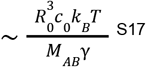, where *γ* is the interfacial tension between the condensate and the buffer phase. In the Cahn-Hilliard formulation, the interfacial tension is defined by the excess free energy per unit area between two phases at equilibrium that form a flat interface. By integrating in the normal direction of the interface from one phase to another,

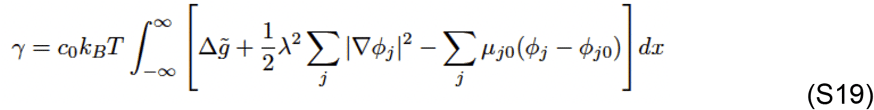

where *µ*_*j*0_ is the chemical potential of component *j, ϕ*_*j*0_ is the volume fraction of component *j* in either phase far away from the interface. It can be shown that *γ* ∼ *λc*_0_*k*_*B*_*T* ^S14,S18^.

Therefore, we define a characteristic diffusion time scale 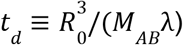. Similarly, the reaction-limited dissolution/growth time scale of the condensate can be derived based on the Allen-Hilliard equation (eliminating the diffusion term in Eq. S11) 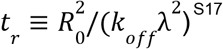. Based on the diffusion and reaction time, we define Damköhler number Da ≡ *t*_*d*_ */ t*_*r*_ =*k*_off_ *R*_0_ *λ / M*_AB_.

The equation of motion of the center of the telomere locus using the Newtonian model motivates us to define another viscously dominated coalescence time *t*_*v,i*_. Because the length scale of the interaction kernel is *λ*, depending on the dimensionality *ν*, the force |**F**_*i*_| ∼ *c*_0_*k*_*B*_*Tλ*^*ν*−1^. Therefore, based on Eq. S15 (*η*_*i*_*R*_0_*/t*_*v,i*_ ∼ |**F**_*i*_|), we define *t*_*vi*_ ≡ *R*_0_*η*_*i*_*/*(*c*_0_*k*_*B*_*Tλ*^*ν*−1^). We may define a dimensionless stiffness constant based on the ratio of the elastic force ∼ *R*_0_*E*_*i*_ and capillary force (|**F**_*i*_| above),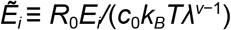. Similarly, for the Kelvin-Voigt and Maxwell models, we define the retardation time *τ*_*i*_ = *η*_*i*_*/E*_*i*_, and the Maxwell relaxation time 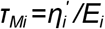. We nondimensionalize these time scales by *t*_*d*_.

We visualize the simulation in images and movies where the red (R), green (G), and blue (B) values are R = B = *ϕ*_C_, G = *ϕ*_A_ + *ϕ*_B_ + *ϕ*_AB_, in other words, magenta is used to denote component C, and green is used to denote the Corelet concentration. The kymographs show the simulation results at the symmetry line connecting the two telomere loci over time.

Here, we provide a summary of the simulations and the corresponding dimensionless parameters.

### Base case simulation

The purpose of the first simulation is to illustrate the possibility that capillary forces can reposition attached objects. A simulation using the dashpot model reproduces the merging of two telomeres pulled by the condensate, shown in Supplemental Figure 3A. Denoting the interfacial tension between the AB-rich (C-rich) phase and the buffer-rich phase by *γ*_AB,S_ and *γ*_C,S_, and that between the AB-rich and C-rich phases by *γ*_AB,C_, then bringing AB-rich and C-rich phases into contact, the affinity between the AB-rich and C-rich phases can be quantified by *γ*_AB,S_ +*γ*_C,S_ −*γ*_AB,C_. In the Cahn-Hilliard formulation, the interfacial tensions defined in Eq. S19 are related to the interaction parameters *Χ*_*ij*_ ^S18^. In Supplemental Figure 3A, *Χ*_AB,S_ = *Χ*_C,S_ = *Χ*_AB,C_ = 2.

Because we are interested in the capillary interaction between the condensate and the chromatin locus, we choose a set of thermodynamic parameters that can give the desired capillary adhesion between components AB and C qualitatively. The precise quantification of the saturation concentration and phase diagram^S12,S19^ is beyond the scope of the modeling here. For the base case simulation where AB is the FUS_N_ Corelet condensate (FUS_N_-mCh-sspB + iLID-GFP-Fe, in activated/interacting state), C is FUS_N_-miRFP670-TRF1, *χ*_AB,S_ = 3, *χ*_C,S_ = 3, *χ*_AB,C_ = 2, *v*_A_ = *v*_B_ = *v*_S_ = 1, *v*_AB_ = *v*_C_ = 2. Suppose when under light activation the association between A and B is strong and *K*_*d*_ = *K*_*d*,on_ = 0, *k* = *k*_on_, while when not illuminated *K*_*d*_ = *K*_*d*,off_ = 0.2, *k* = *k*_off_. Based on the free energy of mixing (Eq. S5) and the volume fraction, the activities of A and B are on the order of 10 ^−2^ while that of AB is on the order of 1. In order for the initial rate of association ∼ *k*_on_*a*_A_*a*_B_ when light is turned on and the initial rate of dissociation ∼ *k*_off_*K*_*d*,off_ *a*_AB_ when light is off to have the same order of magnitude, we set *k*_on_*/k*_off_ = 4 × 10^3^.

We make the assumption that the diffusivity of component C is the same as that of the Corelet condensate component (AB). Based on the measurement of the diffusivities of FUS_N_-mCh-sspB and iLID-GFP-Fe^S12^, we set *M*_A_ = *M*_B_ = 10*M*_AB_ = 10*M*_C_. For the base case, we set the average composition to be *ϕ*_A_+*ϕ*_B_+*ϕ*_AB_ = 0.164, *ϕ*_C_ = 0.041, and suppose the constituents (A and B) of the condensate have the same volume fractions.

For the time scale, we set Da = *t*_*r*_*/t*_*d*_ = 2000; *t*_*v*1_*/t*_*d*_ = *t*_*v*2_*/t*_*d*_ = 0.02, that is, the two telomere loci have the same inverse mobility and are set to a small value so that it does not have a significant effect. Without yet considering the viscoelastic properties, we use the Newtonian model for the base case. Here in this section, the purpose of the simulation is to illustrate the possibility that telomere droplets can be pulled to merge with capillary forces. The ratios of time scales above remain to be studied and validated in later sections given other experimental evidence.

The simulations are performed in a 2D periodic *L* × *L* domain to capture the dynamics qualitatively. For the base case *L/λ* = 36. *R*_0_*/λ* = 9. As an example, in Fig. 1 *R*_0_ ≈ 0.245 µm, which corresponds to *λ* ≈ 27 nm. Initially, the system is fully equilibrated with the presence of two droplets that are rich in component C (which represents the FUS_N_-TRF1-marked telomeres that are observed in microscopy and henceforth called droplet C) and the light is off. With the imposed average composition, the radii of droplet C are 2.5*λ* (defined by the region where *ϕ*_C_ *>* 0.5). Their centers are located at **r**_1_(*t* = 0) = [−*d*_0_*/*2,0], **r**_2_(*t* = 0) = [*d*_0_*/*2,0], where *d*_0_*/λ* = 26. At *t* = 0, a circular region {|**r**| ≤ *d*_0_*/*2} is illuminated. Upon light activation, two Corelet droplets that are rich in AB nucleate in the circular region next to the telomeres. We let the system fully equilibrate until the two droplets merge and form a single condensate in between the two telomere loci. The distance between the two telomere loci becomes 2*R*_0_, where *R*_0_*/λ* = 9, and then turn off the light and let the condensate dissolve. The equations are solved on a grid of [128,128], that is, the grid spacing is 0.3*λ*.

In the following sections where we study the effect of certain parameters, other parameters remain unchanged unless otherwise stated.

### Differential levels of adhesion

Increasing *χ*_AB,C_ is expected to increase the energy of interaction between AB-rich phase and C-rich phase and hence the interfacial tension *γ*_AB,C_, resulting in a decrease in the affinity between AB-rich and C-rich phases, modeling the changes in the condensate and telomere loci. In Supplemental Figures 3C and D, we performed a pair of control simulations where the only difference between them is the value of *χ*_AB,C_. We see that when *χ*_AB,C_ = 2, the adhesion between the telomere and Corelet droplet is strong, the telomeres stay attached to the condensate and eventually merge, whereas when *χ*_AB,C_ = 3.5, the telomeres detach and do not merge. In Supplemental Figure 3B, additional parameters are chosen to match the observed length and time scales in Figure 2A, where there is “no interaction” between the condensate and the chromatin locus. In Supplemental Figure 3A, *χ*_AB,S_ = *χ*_C,S_ = *χ*_AB,C_ = 2, while in Supplemental Figure 3B *χ*_AB,S_ = *χ*_C,S_ = 2, *χ*_AB,C_ = 3.5.

Inspired by experiments shown in Figure 2E, the effect of adhesion is further demonstrated in Supplemental Figure 3F, which shows that when the Corelet droplet is next to a single telomere it stays attached to the telomere during dissolution when *χ*_AB,C_ = 2 and it detaches when *χ*_AB,C_ = 4.7. In Supplemental Figure 6E, we study the effect of viscous resistance using a simulation of the interaction between a single telomere and the condensate. We see that the displacement of the telomere toward the condensate decreases with increasing viscous resistance. The small telomere displacement seen from experiments in Figure 2E in the main text indicates the resistance experienced by the telomere is high compared to the condensate.

We study the effect of changing *χ*_AB,C_ while keeping *χ*_AB,S_ = *χ*_C,S_ = 3 the same as the base case. Compared to Supplemental Figure 3A, in Supplemental Figure 3B we increase *χ*_AB,C_ to 3.5 and keep all parameters identical to the ‘Base case simulation’ section except for the sizes of droplets and the average composition to better reflect the relevant experimental length and time scales: the average compositions are *ϕ*_A_ + *ϕ*_B_ + *ϕ*_AB_ = 0.141, *ϕ*_C_ = 0.038. The distance between the two telomeres initially is *d*_0_*/λ* = 27.3. The illumination region is {|**r**| ≤ *d*_0_*/*2}. When the two Corelet droplets merge into a single droplet, the merged droplet radius is 6.7*λ*. Again, *t*_*r*_ and *t*_*d*_ are still defined using *R*_0_ = 9*λ* as the characteristic length scale. For reasons we will explain in the ’Single telomere interaction with condensate’ section, we increase the viscous resistance of the telomere loci to *t*_*v*1_*/t*_*d*_ = *t*_*v*2_*/t*_*d*_ = 1, and increase the Damköhler number to Da = *t*_*r*_*/t*_*d*_ = 10^4^ in order to slow down droplet dissolution because, at this level of viscous resistance, it becomes easier for the telomere to detach even for the base Flory interaction parameter of *χ*_AB,C_ = 2.

To illustrate the effect of *χ*_AB,C_, we perform a controlled pair of simulations where the only difference is *χ*_AB,C_ shown in Supplemental Figure 3C and D. The average compositions are identical and set to *ϕ*_A_ + *ϕ*_B_ + *ϕ*_AB_ = 0.167, *ϕ*_C_ = 0.038, and the initial distance to *d*_0_*/λ* = 26.7. The illumination region is {|**r**| ≤ *d*_0_*/*2}.

### Single telomere interaction with condensate

In this section, we study the interaction between a single telomere and the condensate. The domain size remains the same as the base case. The average volume fraction of C is increased such that the initial radius of the C-rich phase is *R*_*t*_ = 4.3*λ*, while the average volume fraction of A and B remains the same as the base case. Da = 10^4^ is the same as what is laid out in the ‘Different levels of adhesion between droplets’ section. Initially, the single telomere is placed at [−3.3*λ*, 0]. The illumination region is {|**r**| ≤ *d*_0_*/*2} (where *d*_0_*/λ* = 26, the same as the base case). All other parameters are the same as the base case.

We study the effect of the viscous resistance *η*_1_ by changing *t*_*v*1_*/t*_*d*_, where the subscript 1 refers to the only telomere locus that is considered here. In Supplemental Figure 6E, we show the kymograph of the process of Corelet dissolution (*t* = 0 corresponds to the moment when light is turned off) as a function of *t*_*v*1_*/t*_*d*_. The small telomere displacement seen from experiments in Figure 1E in the main text indicates the resistance to telomere motion is high. This is the reason that in the ‘Different levels of adhesion between droplets’ section we set *t*_*v*1_*/t*_*d*_ = 1.

Next, setting *t*_*v*1_*/t*_*d*_ = 1, we compare simulations with *χ*_AB,C_ = 2 and *χ*_AB,C_ = 4.7 in Supplemental Figure 6F. For both cases, the Corelet condensate nucleates next to the telomere when the light is turned on. During condensate dissolution, when *χ*_AB,C_ = 2, the condensate stays attached to the telomere, due to the stronger adhesion between the two, however, when *χ*_AB,C_ = 4.7, the weak adhesion causes the condensate and telomere to detach from one another.

### Models of chromatin viscoelasticity

In Figure 3 in the main text and Supplemental Figure 3G, we consider models of viscoelasticity including Kelvin-Voigt (Eq. S16) and Jeffreys model (Eq. S18). For better agreement with experiments shown in Figure S2C, namely the distance moved before telomeres detach, we choose normalized retardation time τ_1_ /*t*_*d*_ =τ_2_ /*t*_*d*=_2, and dimensionless resistance 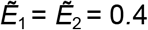, where the subscripts 1 and 2 refer to the two telomere loci. For the comparison between Kelvin-Voigt and Jeffreys models, we vary the normalized Maxwell relaxation time τ_*M,i*_/*t*_*d*_. The Kelvin-Voigt model is a special case of Jeffreys model with τ_*M,i*_=∞.

We do not consider the Maxwell model here because if the impedance on the telomere is described completely by the Maxwell model, then the telomere will exhibit an instantaneous recoil when the condensate detaches. Notice that in our model, the dynamics of the center of telomeric chromatin (**r**_*i*_) and that of TRF1 (component C) are modeled separately, the overall impedance is contributed by both. In our physical model, using a Maxwell model (or a spring model) for the telomeric chromatin will not cause instantaneous recoil, but the recoil dynamics will be faster than a Jeffreys model that has the same spring and dashpot connected in series but with an additional dashpot in parallel. Thus, the retardation time τ/*t*_*d*_ controls the speed of the recoil dynamics and using the Maxwell model is equivalent to setting τ_1_ /*t*_*d*_=τ_2_ /*t*_*d*_=0.

Consistent with the ‘Single telomere interaction with condensate’ section, we set Da = 10^4^. All other parameters in Figure 3 and Supplemental Figure 3G are identical to the base case. All other parameters for the simulations in Supplemental Figures 3-4, except for the ones that are varied as well as the initial conditions, are identical to the base case. The initial condition for Figure 3, Supplemental Figure 3G and S4D-K is the state of the base case at the moment when the light is turned off (at the end of the light-on period).

### Predicting recoil behaviors of detached loci

Because the Newtonian model for telomere loci is unable to predict the recoil when they detach from the condensate observed in experiments, we consider viscoelastic models including the Kelvin-Voigt (Eq. S16) and the Jeffreys models (Eq. S18). Supplemental Figure 3G shows the distance between two telomeres as a function of time at various *t*_*d*_/τ_*M*_ (where τ _*M*_=τ_*M*,1_ =τ_*M*,2_ is the Maxwell relaxation time for both telomere loci). We see that the amount of recoil lessens with decreasing the Maxwell relaxation time. Kymographs predicted by the Kelvin-Voigt model (*t*_*d*_/τ_*M*_=0), which predicts that the telomere recoils back to its original position, and the Jeffreys model (*t*_*d*_/τ _*M*_=0.8), which predicts a partial recoil, are shown in Figure 3 in the main text.

In Supplemental Figure 4, we analyze the asymmetry of viscoelastic properties. We make the assumption that a telomere near the nuclear periphery experiences a proportionately higher spring stiffness and viscous friction, that is, when using the Kelvin-Voigt model, we assume *ρ*_2_*/ρ*_1_ ≡ *E*_2_*/E*_1_ = *η*_2_*/η*_1_, hence the definition of *ρ*_2_*/ρ*_1_ in the main text. Since *t*_*i*_=η_*i*_/*E*_*i*_, this assumption is equivalent to *t*_1_ = *t*_2_.

First, we use the Kelvin-Voigt model, set the retardation time on both sides to be equal τ_1_/*t*_*d*_=τ_2_/*t*_*d*_=2 and fix the normalized stiffness on one side 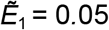 while varying 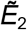 (Supplemental Figure 4D shows the trajectory *x*(*t*) of the two telomeres in this Kelvin-Voigt model). We define Δ*x*_*i*_ to be the distance that the telomere travels from *t* = 0 to the moment when the two telomere merge, defined to be the time point with the highest |*d*^2^ *x*/*d*^2^|, indicated by the filled circles in Supplemental Figure 4D. Supplemental Figure 4E shows that the ratio Δ*x*_1_*/*Δ*x*_2_ increases linearly with increasing 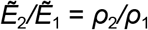. Considering the resistance of the condensate in between the two telomeres, we use a simple model to predict the asymmetry in the displacement given the asymmetry in the viscoelastic properties,

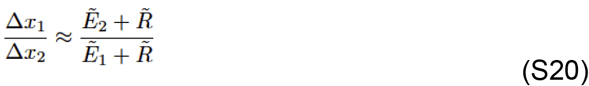

where 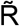 is the effective resistance of the condensate relative to that of the telomeric chromatin. Fitting the data in Supplemental Figure 4E to Eq. S20, we obtain 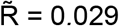.

Next, we fix the ratio of the stiffness constants 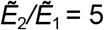 and vary 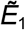. Supplemental Figure 4F shows the trajectories and S4G shows that the ratio Δ*x*_1_*/*Δ*x*_2_ gradually increases and reaches a plateau with increasing 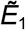. Therefore, when the resistance from the chromatin dominates over that of the condensate, which is likely the case given the negligible displacement of the telomere when a single telomere is attached to the Corelet droplet as it dissolves, the ratio of displacement becomes closer to the inverse ratio of the stiffness 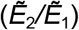. Fitting the data to Eq. S20, we obtain 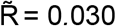.

In Supplemental Figure 4H-K, we repeat the analysis above using the Jeffreys model. Again, we assume that all dashpot and spring elements have proportionately higher values near the nuclear periphery, that is, 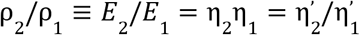. In other words, both telomeres have the same retardation time and Maxwell relaxation time. We set *τ*_1_*/t*_*d*_ = *τ*_2_*/t*_*d*_ = 2 and *τ*_*M*,1_*/t*_*d*_ = *τ*_*M*,2_*/t*_*d*_ = 2. Again, we find that Eq. S20 is a good approximation for the ratio of displacement and the fitted value of 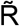 is 0.042 and 0.041 in Supplemental Figure 4I and K, respectively.

The consistency in the fitted 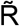 indicates that Eq. S20 provides a useful estimate of the asymmetry. More importantly, Eq. S20 provides an upper bound to the ratio of the displacement equation

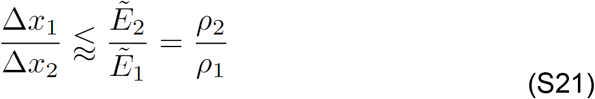

In other words, given the ratio of displacement observed in experiments, Eq. S21 provides a lower bound for the ratio of local resistance.

### Repositioning non-chromatin nuclear bodies

A simulation of the Cajal body in Figure 5 in the main text and Supplemental Figure 5D uses Eq. S6 without the interaction kernel. Parameters are chosen to match the observed length and time scales in Figure 5D. We also perform an additional control simulation comparing the telomere (using Eq. S13) and the Cajal body Eq. S6 with the only difference being whether the interaction kernel is included. The comparison in Figure 5E and Supplemental Figure 5D and E shows that the coilin construct forms a ring at the interface between the condensate and the buffer, while FUS_N_-TRF1 does not form a ring around the Corelet droplet due to its attractive interaction with telomere loci.

The simulation shown in Figure 5 and Supplemental Figure 5D-E reproduces the experimental observation that when FUS_N_ is fused with coilin rather than TRF1, Cajal bodies can also be merged by the Corelet droplet via capillary forces. The relevant difference between modeling telomeres and Cajal bodies in this simulation is that the Coilin protein is not directly chromatin bound. Therefore, we remove the interaction kernel and use only the Flory-Huggins chemical free energy of mixing to represent a Cajal body. Here, we perform simulations with all the parameters identical to the ‘Base case simulation’ section except for the sizes of the condensates and domain and the average composition to better reflect the experimental observation in Figure 5D. The average compositions are *ϕ*_A_ + *ϕ*_B_ + *ϕ*_AB_ = 0.104, *ϕ*_C_ = 0.03 The domain size is changed to *L/λ* = 50. Initially, the radii of the two Cajal bodies are 2.3*λ*. Their centers are located in the same way as the ‘Base case simulation’ section, while the distance is *d*_0_*/λ* = 33. The illumination region is again {|**r**| ≤ *d*_0_*/*2}. When the two condensates merge into a single droplet, the radius of the merged droplet is 8*λ*. Note that while the single condensate size is different from *R*_0_ in the ‘Base case simulation’ section, *t*_*r*_ and *t*_*d*_ are still defined using *R*_0_ = 9*λ* as the characteristic length scale.

In Supplemental Figures 5D and E, we compare the telomere (with interaction kernel) and Cajal bodies (without interaction kernel) using the same exact parameters, average composition, and length scales as defined above.

### Estimating number of IDR-IDR interactions at the chromatin-condensate interface

Previously, we have measured the diameter of telomeres in U2OS cells by super-resolution STED microscopy to be on the order of 100-150 nm (data not shown), in agreement with published estimates of telomere sizes in different cell lines^S3-S8^. The surface area of an average telomere would then be 4*π*(0.075µm)^2^ = 0.071µm^2^. We estimate from our images that usually less than half of the surface of the telomere is in contact with the Corelet condensate, so the surface area of the chromatin-condensate interface is up to 0.035 µm^2^. Each FUS_N_-sspB-decorated ferritin “core” particle has a radius of 20 nm^S12^, so a surface area of 0.005µm^2^. Assuming packed spherical core particles coating the condensate-telomere interface, we estimate that approximately (0.035/0.0025) = 14 cores interact with the available surface of the telomere. Every core has binding sites up to 24 FUS_N_ IDRs distributed on its surface, up to half of which (12) are in an orientation to interact with the locus-tethered IDRs at any time. These experiments represent a bulk of approximately 14 × 12 = 168 condensate IDRs interacting with a similar number of chromatin-tethered IDRs providing the interfacial force between a chromatin locus and condensate.

### Limitations of preceding automated activation protocols

First, we attempted a global activation pattern across the entire nucleus, which results in Corelet condensates nucleating at each telomere, but infrequent condensate coalescence events due to small overall condensate size (Figure 5A). The smaller an area of the nucleus that is activated, the larger each condensate grows; so in our next attempt we activated a smaller rectangular region, then shifted the activation region of interest (ROI) across the nucleus over time to activate the nuclear area sequentially (Figure 5A). This sliding box pattern resulted in larger condensates that do fuse, but is prohibitively time-consuming at 60 minutes per nucleus. Next we tested an array pattern of activation sites, reasoning that some of the array positions will nucleate between closely positioned telomeres and lead to coalescence events (Figure 5A); while this pattern does sporadically create condensates at productive locations, most are not aligned with telomere loci positions, and thus its efficiency in merging multiple condensates associated with telomeres is low. The best approach for successful repositioning was to identify close telomere pairs at most 1.4 microns apart using a feedback protocol and to only activate those identified pairs, forming condensates at those positions (Figure 5B).

